# Cavin1 intrinsically disordered domains are essential for fuzzy electrostatic interactions and caveola formation

**DOI:** 10.1101/831149

**Authors:** Vikas A. Tillu, James Rae, Ya Gao, Nicholas Ariotti, Matthias Floetenmeyer, Oleksiy Kovtun, Kerrie-Ann McMahon, Natasha Chaudhary, Robert G. Parton, Brett M. Collins

**Author notes:** Corresponding authors: Robert Parton, Brett Collins.

## Abstract

Caveolae are spherically shaped nanodomains of the plasma membrane, generated by cooperative assembly of caveolin and cavin proteins. Cavins are cytosolic peripheral membrane proteins with negatively charged intrinsically disordered regions (DR1-3) that flank positively charged *α*-helical regions (HR1 and HR2). Here we show that the three DR domains of Cavin1 are essential for caveola formation and dynamic trafficking of caveolae. Electrostatic interactions between DR and HR regions promote liquid-liquid phase separation behaviour of Cavin1 *in vitro*, assembly of Cavin1 oligomers in solution, generation of membrane curvature, association with caveolin-1 (CAV1), and Cavin1 recruitment to caveolae in cells. Removal of the first disordered region causes irreversible gel formation *in vitro* and results in aberrant caveola trafficking through the endosomal system. We propose a model for caveola assembly whereby fuzzy electrostatic interactions between Cavin1 and CAV1 proteins, combined with membrane lipid interactions, are required to generate membrane curvature and a metastable caveola coat.

## Introduction

Caveolae (‘*little caves*’) are membrane invaginations with a diameter of 50-60 nm that are abundant in the plasma membrane of many cell types such as muscle fibres, endothelial cells and adipocytes. These membrane nanodomains are important for an array of different functions including endocytosis, intracellular signalling, lipid and fatty acid homeostasis and response to membrane stress 1-3.

Although the precise details of caveola biogenesis remain enigmatic their assembly requires the activities of two families of proteins - caveolins and cavins - and their coordinated interactions with membrane lipids and cholesterol. The integral membrane proteins of the caveolin family (CAV1, CAV2 and muscle specific CAV3) are synthesized at the endoplasmic reticulum and trafficked via the Golgi apparatus to the plasma membrane ^4^. Caveolins have an unusual hairpin structure that inserts into the membrane bilayer, with an extended N-terminal domain and *α*-helical C-terminal domain exposed to the cytoplasm ^3, 5, 6^. When expressed on its own in mammalian cells the core caveolin CAV1 is diffusely localised in the plasma membrane and is unable to form spherical caveolae in the absence of cavins ^4, 7^. In contrast, CAV1 is able to generate membrane vesicles similar to caveolae (h-caveolae) upon heterologous expression in *Escherichia coli* ^8^. This points to an intrinsic capacity of CAV1 to generate membrane curvature, which is thought to be enabled by the specific lipid composition of *E. coli* membranes. In metazoan cells however, the additional presence of the peripheral membrane cavin proteins is required for the formation of native caveolae. In particular, Cavin1 and CAV1 are together required and sufficient to generate a minimal core system for caveola formation at the plasma membrane. Other cavin family members require Cavin1 for their recruitment and are thought to provide regulatory or tissue-specific activities ^7, 9, 10^.

All cavin proteins share a highly characteristic domain architecture consisting of two core *α*-helical regions (HR1 and HR2) with relatively high sequence conservation ^11, 12^. These are connected by three intrinsically disordered regions (DR1, DR2 and DR3), that possess very little sequence homology but share the property of being enriched in negatively charged residues (**Fig. 1A**). Cavin proteins can assemble into homo- and hetero-oligomeric complexes that form a protein coat on the cytosolic face of caveolae; and the essential isoform Cavin1 can form homo-oligomers that drive caveola formation in the absence of other family members ^11, 13, 14^. The N-terminal *α*-helical HR1 domain of Cavin1 forms a core trimeric coiled-coil structure that also promotes heteromeric interactions between other members of the Cavin family ^11^. A surface exposed patch of basic amino acid residues in the HR1 domain has affinity for phosphoinositide lipid headgroups including phosphatidylinositol-4,5-bisphosphate (PI(4,5)*P*_2_) ^11^. The C-terminal *α*-helical HR2 region of Cavin1 is unique in the cavin family as it also contains a stretch of repeated undecad sequences (11-mers) predicted to form a second coiled-coil structure termed UC1 (undecad of Cavin1) ^10^. Basic amino acids within the HR2 and UC1 domains can associate with phosphatidylserine (PS) to regulate caveola formation and stability ^10^. These two *α*-helical lipid interacting sites are important for membrane recruitment and for generating caveolar membrane curvature. However, the molecular mechanisms of caveolar membrane association and higher-order assembly of cavins with caveolins at the cell surface are largely unknown.

**Figure 1.**
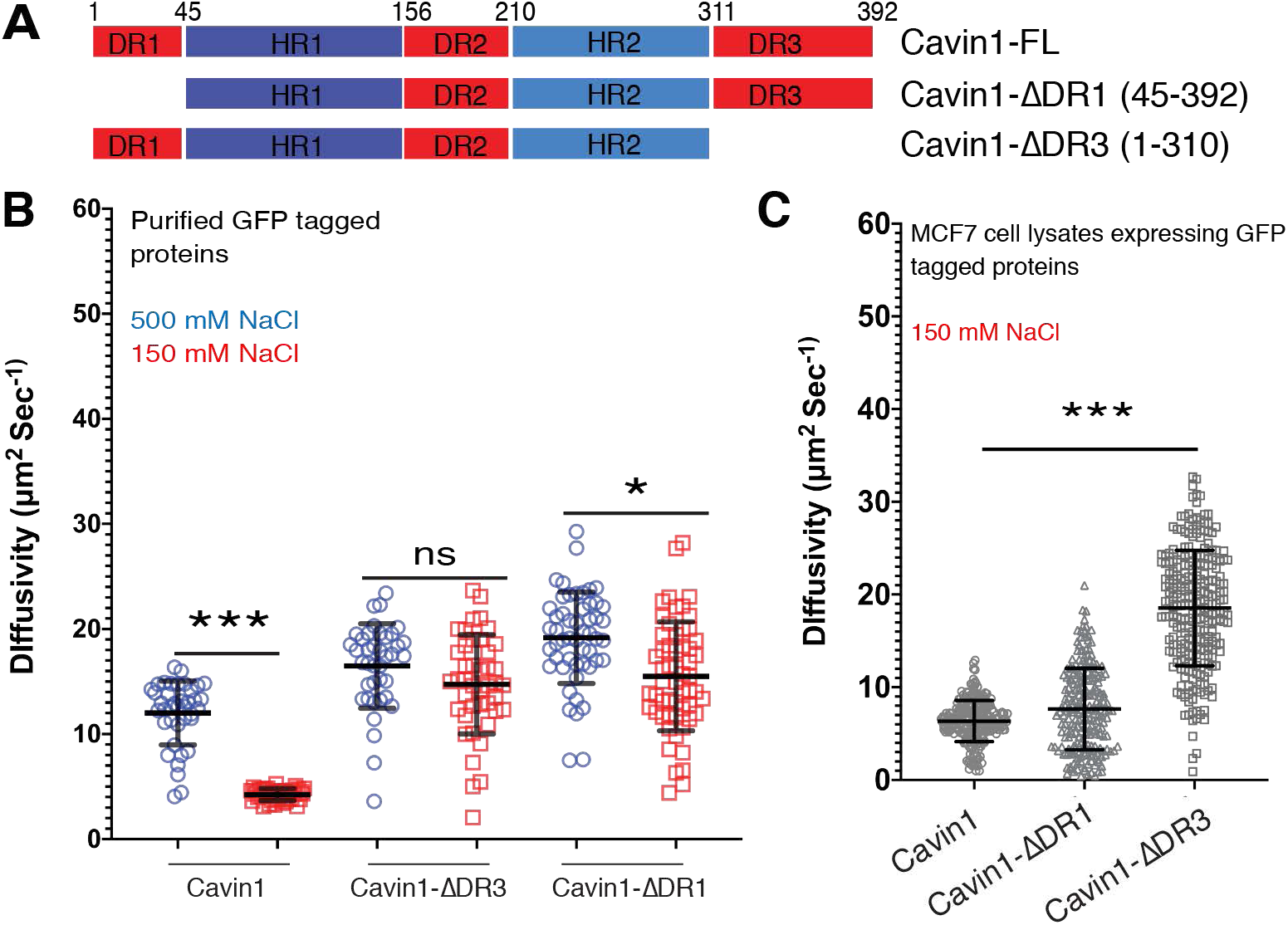
The Cavin1 N- and C-terminal DR domains are important for self-association into oligomers. **(A)** Schematic representation of Cavin1 and truncations. DR, disordered region; HR, helical region. **(B)** The diffusion rate of Cavin1, Cavin1-ΔDR1 and Cavin1-ΔDR3 in solution assessed by fluorescence correlation spectroscopy (FCS). Bacterially expressed and purified ubiquitin and GFP tagged proteins (**Fig. S2**) were analysed in high NaCl concentration (500 mM) and physiological NaCl concentration (150 mM). Error bars indicate mean ± SD (standard deviation), N=2, n=10-15, ns – not significant, *P<0.05 ***P<0.001. **(C)** The diffusion rate of GFP-tagged Cavin1, Cavin1-ΔDR1 and Cavin1-ΔDR3 in lysates after expression in MCF7 cells (lacking endogenous Cavins and Caveolins). Buffer contained 150 mM NaCl. N=3, n=20-25, ns – not significant, ***P<0.001. Error bars indicate mean ± SD.

In this study we examined the role of the uncharacterised DR domains of Cavin1 in caveola formation. The DR domains of Cavin1 are strictly required for caveola assembly, and a systematic dissection of these intrinsically disordered regions showed that there are minimal acidic sequences within the DR domains that are essential for caveolar targeting, *in vitro* membrane remodelling and homo-oligomeric Cavin1 complex assembly. We find that Cavin1 undergoes electrostatically driven self-association via its disordered regions that promotes liquid-liquid phase separation (LLPS) *in vitro*, that it can co-phase separate with CAV1, and this is dependent on specific sequence properties of the two proteins. Perturbing the DR domain-mediated dynamics of Cavin1 self-association has profound effects on Cavin1 and CAV1 localisation and caveolar trafficking in cells. Our results lead us to propose a model for caveola assembly involving ‘fuzzy’ electrostatic interactions by Cavin1 at the CAV1/membrane interface to generate a metastable caveola coat.

## Results

### Cavin1 forms electrostatically driven oligomers that depend on DR1 and DR3 domains

The cavin family proteins all share distinguishing structural similarities with each other, consisting of disordered N- and C-terminal domains DR1 and DR3, a central disordered region DR2, and interspersed *α*-helical coiled-coil region HR1 and predicted *α*-helical region HR2 (**Fig. 1A**). The trimeric coiled-coil HR1 domain and C-terminal HR2 domains are both rich in basic amino acid residues, while the three DR domains instead possess a high proportion of acidic amino acid residues. This alternating electrostatic charge distribution is a distinctive and conserved feature of all family members (**Fig. S1A**), indicating it is an essential characteristic of the proteins. We also used the D2P2 web server ^15^ to analyse the sequence of Cavin1 for predicted regions of disorder, and confirmed that the DR1, DR2 and DR3 regions are predicted to be intrinsically disordered as suggested by previous secondary structure analyses ^10, 11^ (**Fig. S1B**). Interestingly, sites of phosphorylation in Cavin1 are predominantly found in the DR1, DR2 and DR3 domains, while sites of ubiquitylation are concentrated in the HR1 and HR2 regions. In subsequent experiments the boundaries of the mouse Cavin1 domains are defined as: DR1 (1-44), HR1 (45-155), DR2 (156-209), HR2 (210-310), and DR3 (311-392) (**Fig. 1A**). Expression constructs used in this study are outlined in **Fig. S2**.

We recently proposed that the predominantly negatively charged DR sequences of Cavin1 may associate with the positively charged HR domains to promote intra and/or inter molecular electrostatic interactions required for coat assembly ^12^. To probe the role of electrostatic interactions, we used fluorescence correlation spectroscopy (FCS) to measure the diffusional properties of purified GFP-tagged Cavin1 in both 500 mM NaCl (high salt) and 150 mM NaCl (iso-osmotic salt concentration). According to polymer theory, the diffusivity of protein molecules in solution decreases with increasing intermolecular interactions due to molecular crowding limiting its molecular motion ^16^. GFP-Cavin1 (100 nM concentration) showed a remarkable decrease in its diffusivity with the reduction of ionic strength from 500 mM NaCl (12.01 ± 3.03 µm^2^/sec) to 150 mM NaCl (4.02 ± 0.50 µm^2^/sec) (**Fig. 1B**). This indicates that at physiological salt concentrations Cavin1 can form homomeric oligomers with an average hydrodynamic radius ∼55 nm, similar to those observed previously ^4, 14^, and that this self-association is dependent on electrostatic interactions. In contrast to full-length Cavin1, removal of either N- or C-terminal DR1 or DR3 domains prevents this electrostatically driven self-assembly at lower physiological salt concentrations (**Fig. 1B**).

Next, we sought to understand the role of DR sequences in oligomeric assembly of Cavin1 in a more representative cellular milieu. For these experiments we used MCF7 cells, which lack caveolae and do not express any caveolin or cavin proteins ^14, 17, 18^. GFP-tagged Cavin1 proteins were transiently expressed and FCS analysis was used to measure the diffusivity of each protein in cell lysates (all at 150 mM NaCl). Full length GFP-Cavin1 in MCF7 cell lysates forms relatively heterogenous large molecular weight species in solution with slow diffusive properties (6.35 ± 2.35 µm^2^/sec) (**Fig. 1C**) similar to purified recombinant GFP-Cavin1. In contrast to purified recombinant GFP-Cavin1-ΔDR1, the N-terminal DR1 deletion in cell lysates showed a similar (although tending to faster) rate of diffusion to the full-length protein (7.65 ± 4.40 µm^2^/sec) (**Fig. 1C**). Complete deletion of the C-terminal DR3 region of GFP-Cavin1-ΔDR3 however, significantly increased the diffusivity of Cavin1 in MCF7 lysates (18.54 ± 6.22 µm^2^/sec), similar to the recombinant GFP-Cavin1-ΔDR3 (**Fig. 1C**). Overall these studies demonstrate a role for the DR sequences in electrostatically driven oligomerisation of Cavin1 in solution.

### Cavin1 undergoes liquid-liquid phase separation (LLPS) influenced by the DR domains

There is an increasing awareness of the role of intrinsically disordered sequences in generating membraneless organelles via liquid-liquid phase separation (LLPS) or ‘demixing’ of proteins and associated molecules in solution. Demixing or LLPS can be driven by a variety of mechanisms, including cation-*π* and *π*-*π* stacking, interactions with polyanions such as RNA, and intermolecular electrostatic interactions ^19–22^. A number of recent studies have shown that membranes can be platforms for nucleating and transporting phase-separated assemblies, or in turn be regulated and organised via LLPS-mediated processes ^23–34^. It has also been proposed that the formation of phase-separated condensates can perform physical work on their surroundings, including at the membrane-cytosol interface to generate membrane curvature ^35–38^. Because of the demonstrated importance of disordered regions in Cavin1 for its assembly behaviour, we assessed whether purified Cavin1 is able to form supramolecular assemblies leading to LLPS *in vitro*.

Purified recombinant GFP-Cavin1 expressed in *E. coli* remains dispersed in solution at both physiological NaCl concentration (150 mM) and at high NaCl concentration (750 mM) within a protein concentration range of 1 to 10 µM (**Fig. S3A**). However, when Dextran T-500 (1.25% w/v) was added as a macromolecular crowding agent ^39, 40^ full-length GFP-Cavin1 rapidly formed spherical liquid droplets at 150 mM NaCl even at low protein concentrations (0.1 µM) (**Fig. 2A; Fig. S3B and Fig. S3C**). This is well below the estimated cellular concentration of Cavin1 of 3 μM ^41^. These droplets increased in size with increasing protein concentration in the range 1 to 10 µM (**Fig. 2A**). Increasing the salt concentration strongly inhibited the ability of GFP-Cavin1 to undergo LLPS, consistent with a role for electrostatic intermolecular interactions ^20, 42^. We also tested both purified full-length Cavin1-GFP isolated from mammalian HEK293 cells, and unpurified Cavin1-GFP in MCF7 cell lysates, and found that both preparations underwent similar salt-sensitive LLPS (**Fig. S3D and S3E**) although this required a higher concentration of Dextran T-500 (3%), possibly due to the presence of other bound proteins, lipids, or post-translational modifications such as phosphorylation, ubiquitylation and SUMOylation present in the mammalian cell expression system partially modifying the properties of Cavin1. Higher concentrations of Dextran T-500 (3%) did not significantly alter the LLPS behaviour of GFP-Cavin1 (**Fig. S3F**). Lastly, we assessed if Cavin1 could undergo LLPS in cells. When over-expressed in MCF7 cells, which lack caveolae due to the absence of CAV1, we found that GFP-Cavin1 on its own remained diffuse and did not form droplets (**Fig. S4A and S4B**). However, after testing several conditions we discovered that if cells were treated with cholesterol following serum starvation GFP-Cavin1 rapidly formed cytoplasmic condensates as well as membrane-associated tubules (**Fig. S4C and S4D**). We then expressed GFP-Cavin1 in MCF7 cells together with an mCherry-CAAX construct as a plasma membrane marker (**Fig. S4E**). After cholesterol addition, we observed plasma membrane localization of GFP-Cavin1 and formation of GFP-Cavin1 and mCherry-CAAX positive plasma membrane-associated tubules. Similar results were observed in CAV1-/-mouse embryonic fibroblasts (MEFs) (**Fig. S4F**). We speculate that cholesterol may alter the normal equilibrium of Cavin1’s interaction with phospholipid membranes, thus promoting self-association and condensation.

**Figure 2.**
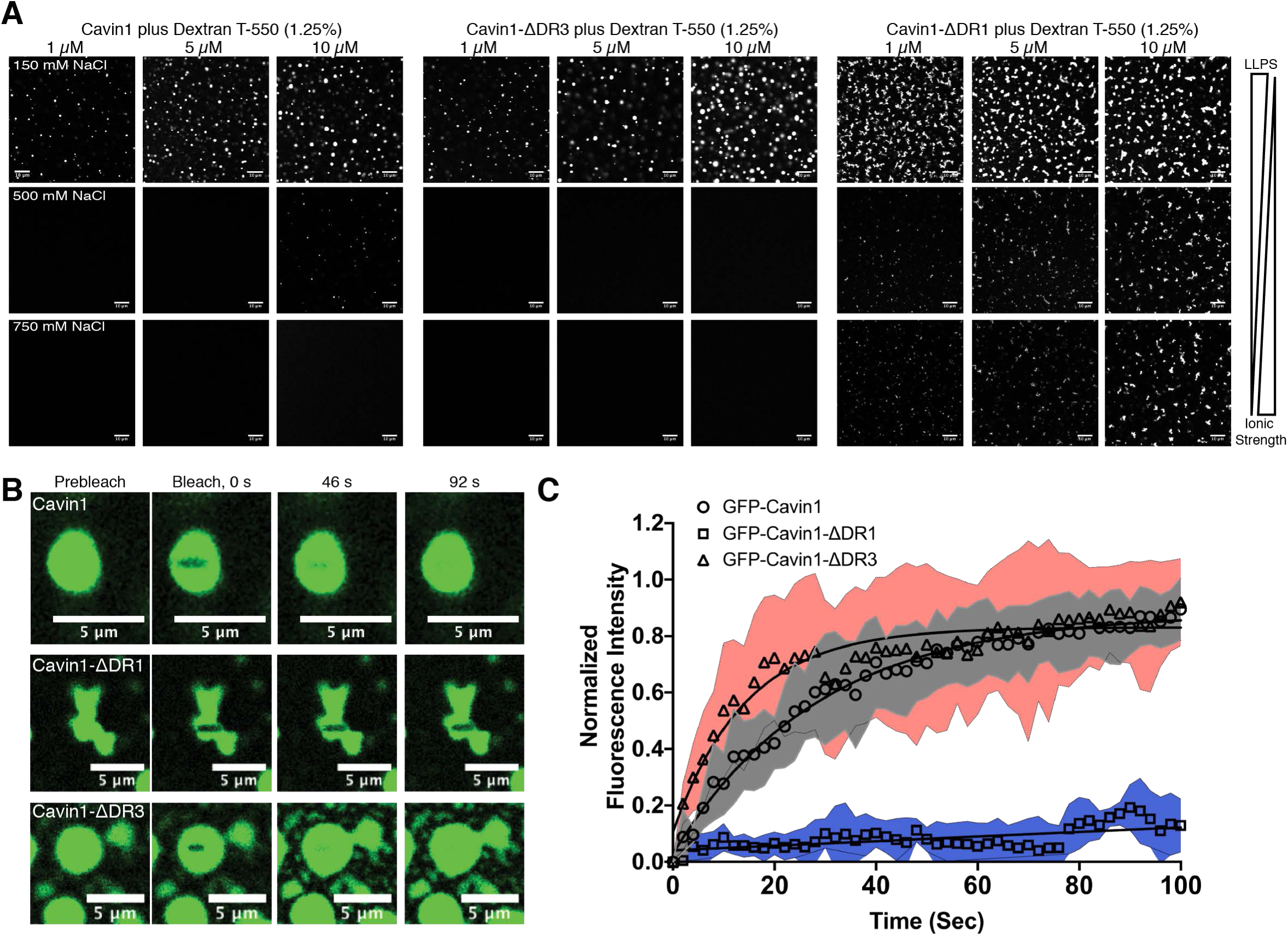
Cavin1 undergoes liquid-liquid phase separation *in vitro*. **(A)** Liquid-liquid phase separation (LLPS) assays with recombinant Ub- and GFP-tagged Cavin1, Cavin1-ΔDR3 and Cavin1-ΔDR1 at different protein and salt concentrations. Scale bar = 10 μm. **(B)** Fluorescence recovery after photobleaching (FRAP) assay with Cavin1, Cavin1-ΔDR3 and Cavin1-ΔDR1 showing GFP fluorescence images at increasing times. Scale bar = 5 μm**. (C)** Plot of normalized fluorescence intensity after photobleaching. N=6-8, Grey, blue and pink shaded areas around recovery curves represent standard deviation (SD). While Cavin1 and Cavin1-ΔDR3 droplets rapidly exchange with the bulk solution and recover their fluorescence, Cavin1-ΔDR1 shows virtually no exchange indicating gel formation.

Deletion of the C-terminal DR3 domain had a small but reproducible effect on the tendency of Cavin1 to undergo LLPS *in vitro*, with droplet formation showing greater sensitivity to increasing ionic strength and protein concentration (**Fig. 2A; Fig. S3B and S3C**). Deletion of the DR1 domain however had a dramatic effect, leading Cavin1 to transition into non-spherical coacervates at all protein and salt concentrations (**Fig. 2A; Fig. S3B and S3C**). Fluorescence recovery after photobleaching (FRAP) was used to analyse the diffusion of proteins within the liquid droplets and the ability of GFP-Cavin1 to exchange with bulk solution. GFP-Cavin1 (*τ*1/2 ∼20s) and GFP-Cavin1-ΔDR3 (*τ*1/2 ∼10s) showed rapid fluorescence recovery after photobleaching, indicating there is ready exchange of protein molecules within the droplets as expected for liquid droplets **(Fig. 2B, 2C**). GFP-Cavin1-ΔDR1, however, showed virtually no recovery **(Fig. 2B, 2C**), suggesting that gel formation has occurred and the truncated protein is unable to diffuse within the condensates ^19, 43^. Overall, these analyses highlight the importance of electrostatic interactions in promoting self-association and subsequent LLPS behaviour by Cavin1, and points to distinct roles of DR1 and DR3 sequences in this process.

### Cavin1 promotes co-phase separation with N-terminal regions of CAV1

Although Cavin1 and CAV1 are associated together in caveolae, it remains unclear whether they interact with each other via direct protein-protein interactions. CAV1 has a unique structural domain architecture shared with other caveolins, consisting of an N-terminal disordered region (DR) (1-60), followed by an oligomerization domain (OD) (61-80), scaffolding domain (CSD) (81-100), intramembrane domain (IMD) (101-133) and a C-terminal membrane associated *α*-helical domain (134-179) (**Fig. 3A; Fig. S5A and S5B**) ^2, 5^. We hypothesized that the N-terminal disordered sequence of CAV1 may enable CAV1-Cavin1 association through interactions involving liquid-liquid phase separation. To test this, we first purified full length CAV1 fused with Maltose binding protein (MBP) and GFP-binding nanobody protein (GBP) ^44^ in non-ionic detergent n-dodecyl β-D-maltoside (DDM) (1.2 mM). MBP-GBP-CAV1 was labelled bound with GFP for visualization, and unlabelled purified Cavin1 was used in a co-phase separation assay (**Fig. 3B and 3C**). Like GFP-Cavin1, unlabelled Cavin1 formed liquid droplets with addition of dextran T-500 (1.25% w/v) observed in bright field image as transparent liquid drops. In the absence of Cavin1 MBP-GBP-CAV1 did not undergo LLPS on its own (**Fig. 3B**). However, when GFP-labelled MBP-GBP-CAV1 was mixed with Cavin1 it was recruited to Cavin1 liquid droplets (**Fig. 3C**). Interestingly MBP-GBP-CAV1 appeared to form a shell around the Cavin1 droplets rather than complete co-mixing.

**Figure 3.**
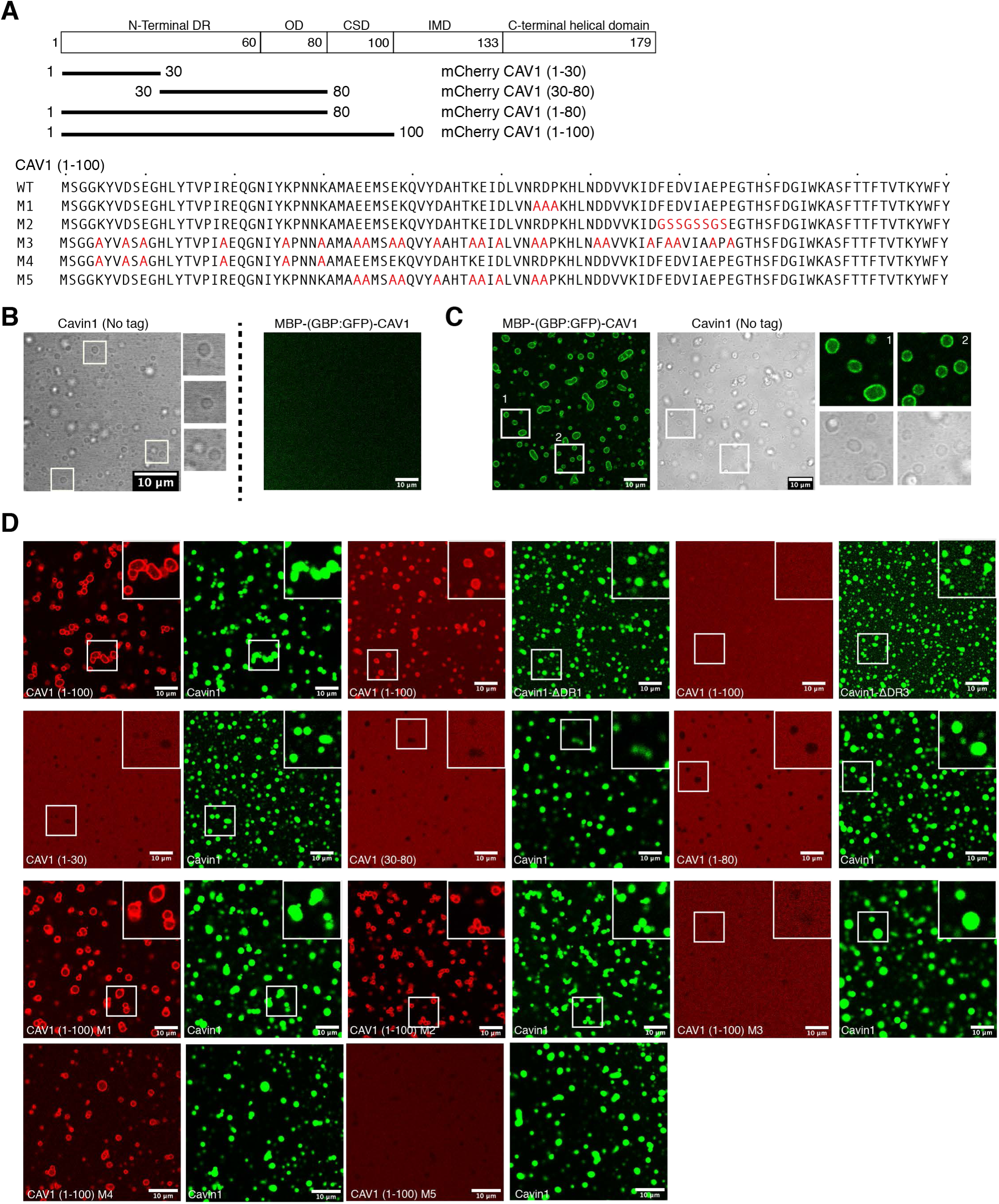
CAV1 N-terminus co-phase separates with Cavin1. (**A**) Schematic representation of CAV1 domain architecture and design of various domain / point mutations. LLPS assays with MBP-GBP-CAV1 and cavin1 independently (**B**) and in mixture (**C**). MBP-GBP-CAV1 does not undergo LLPS in isolation but co-phase separates with Cavin1. (**D**) LLPS assays with different CAV1 DR region mutations and GFP-Cavin1 or Cavin1-DDR3 or Cavin1-DR1. Scale bar – 10 μm. Among all truncation mutations tested, only mCherry-CAV1 (1-100) was able to co-phase separate with Cavin1. Among CAV1 DR point mutations (M1 to M5), mutants M1, M2 and M4 were able to co-phase separate with GFP-Cavin1 while total charge inversion mutant M3 and mutant M5 failed to co-phase separate highlighting the importance of charged residues in CAV1-Cavin1 association.

We next probed the mutual roles of Cavin1 and CAV1 disordered sequences in Cavin1-CAV1 co-phase separation. To this end we generated several mCherry-tagged truncation mutants in the region CAV1 (1-100) encompassing the disordered N-terminus, oligomerization and scaffolding domains (**Fig. 3A**). Similar to previous reports we found that mCherry-CAV1(1-100) formed a higher molecular weight oligomer by gel-filtration, while any truncations of this sequence resulted in monomeric proteins (**Fig. S5C**) ^13^. Like full-length MBP-GBP-CAV1, the mCherry-CAV1(1-100) sequence was able to undergo LLPS with GFP-Cavin1 droplets, again forming an outer shell around the core GFP-Cavin1 droplets (**Fig. 3D**). In contrast truncated mCherry-CAV1 constructs (1-30), (30-80) and (1-80) were all unable to co-phase separate with GFP-Cavin1, and we also observed similar results using unlabelled Cavin1 (**Fig. S5D**). Interestingly, while mCherry-CAV1(1-100) was able to co-phase separate with the Cavin1 N-terminal deletion GFP-Cavin1-ΔDR1, it did not associate with droplets formed by the C-terminal deletion of GFP-Cavin1-ΔDR3 (**Fig. 3D**) suggesting that the Cavin1 DR3 sequences are essential for CAV1 – Cavin1 association.

Sequence alignment of CAV1, CAV2 and CAV3 highlighted several interesting features including an overall conserved but disordered region (30-80) containing two identical motifs, ^54^RDP^56^ and ^68^FEDVIAEP^75^ (**Fig. S5A and S5B**). The N-terminal CAV1 disordered region (1-30) however, was not conserved in CAV2 or CAV3. To further pinpoint the sequence requirements of CAV1 and Cavin1 interaction, we made five mutations in mCherry-CAV1(1-100) (**Fig. 3A**). The two highly conserved motifs ^54^RDP^56^ and ^68^FEDVIAEP^75^ were mutated to alanine, or random glycine and serine (mutants M1 and M2 respectively). The last three mutants (M3, M4 and M5) replaced charged residues (Glu, Asp, Arg, Lys) with alanine in the entire disordered region (1-80) (mutant M3), non-conserved DR fragment (1-30) (mutant M4) and conserved DR fragment (30-60) (mutant M5). CAV1 mutants M1, M2, M4 and M5 all formed oligomers similar to wild-type mCherry-CAV1(1-100) as assessed by their gel filtration profiles, whereas mutant M3 surprisingly migrated as a monomer (**Fig. S5C**). Mutants mCherry-CAV1(1-100) M1, M2 and M4 underwent co-phase separation with GFP-Cavin1 similar to the wild-type CAV1(1-100), while mutants M3 and M5 failed to associate with Cavin1 droplets (**Fig. 3D**). These results confirm that the mCherry-CAV1(1-100) interaction with GFP-Cavin1 is highly specific and depends on charged residues within the CAV1(30-60) region. Overall, these studies indicate that the association between CAV1 and Cavin1 may be driven at least in part by interactions involving liquid phase condensation, with co-mixing mediated by their respective disordered sequences.

Studies of caveolin mutants in cells are typically challenging due to their disrupted trafficking and mis-localisation ^2, 4, 6, 45–51^. Nevertheless, we assessed the localisation of the N-terminal GFP tagged CAV1 mutants (M1 – M5) in the context of the full-length protein and in the presence of Cavin1-mCherry in MCF7 cells (**Fig. S6A**). GFP-CAV1-WT showed the familiar punctate distribution in MCF7 cells that co-localised with Cavin1-mCherry, as did the mutant M4. In contrast the GFP-CAV1 mutants M1 and M5 were not associated with mCherry-Cavin1 at the plasma membrane, and the M2 and M3 mutants were either not expressed or rapidly degraded and could not be detected. Comparison with several organelle markers indicated that mutant GFP-CAV1-M1 was mis-trafficked and accumulated in the Golgi, similar to what was seen with the analogous CAV3(R26Q) dystrophic mutation ^45^ (**Figs. S6B-S6E**). GFP-CAV1-M5 was mostly mis-localised to lipid droplets, with some diffuse plasma membrane localisation and overlap with endosomes. This phenotype was similar to that observed previously when a putative COPII-binding sequence in the CAV1 N-terminus was mutated (D67G) ^4^. Overall these experiments generally correlate with *in vitro* studies, where mutations impacting co-phase separation with Cavin1 do not associate with Cavin1 at the cell surface, are unable to form caveolae and are either degraded or mis-localised in cells.

### The Cavin1 DR sequences are essential for membrane remodelling *in vitro*

We previously showed that Cavin1 and Cavin2 possess an intrinsic ability to tubulate artificial lipid membranes using negative stain electron microscopy ^11^. To examine this membrane remodelling by Cavin1 at higher resolution, we first performed cryoelectron microscopy (cryoEM) analysis of samples after mixing purified Cavin1 with small unilamellar vesicles (SUVs) composed of Folch lipid extracts. We observed formation of an extensive network of membrane tubules (34 ± 5 nm, 12 tubules, 2 independent experiments) possessing a Cavin1 protein coat using both negative stain electron microscopy and cryoEM (**Fig. 4A**). Although tubulation of Folch membranes was most efficient, Cavin1 could also tubulate liposomes consisting of PC/PE/PI(4,5)*P*_2_ (**Fig. S7A**). In addition, Cavin1-GFP expressed and purified from HEK293 cells could also tubulate Folch membranes similarly to the bacterially expressed protein (**Fig. S7B**). Examination of the Cavin1-coated tubules by cryoelectron tomography (cryoET) revealed a striated but relatively heterogeneous pattern of protein densities around the tubules (**Fig. 4B; Movie S1**). These are similar to structures previously observed on the cytosolic face of caveolae using fast-freeze deep-etch ^52, 53^ and conventional EM methods ^54–56^, and with the elongated rod-like structures of isolated Cavin1 observed by negative staining EM ^11^. These experiments indicate that Cavin1 possesses an inherent membrane remodelling activity, driven by large scale oligomeric assembly on the membrane surface.

**Figure 4.**
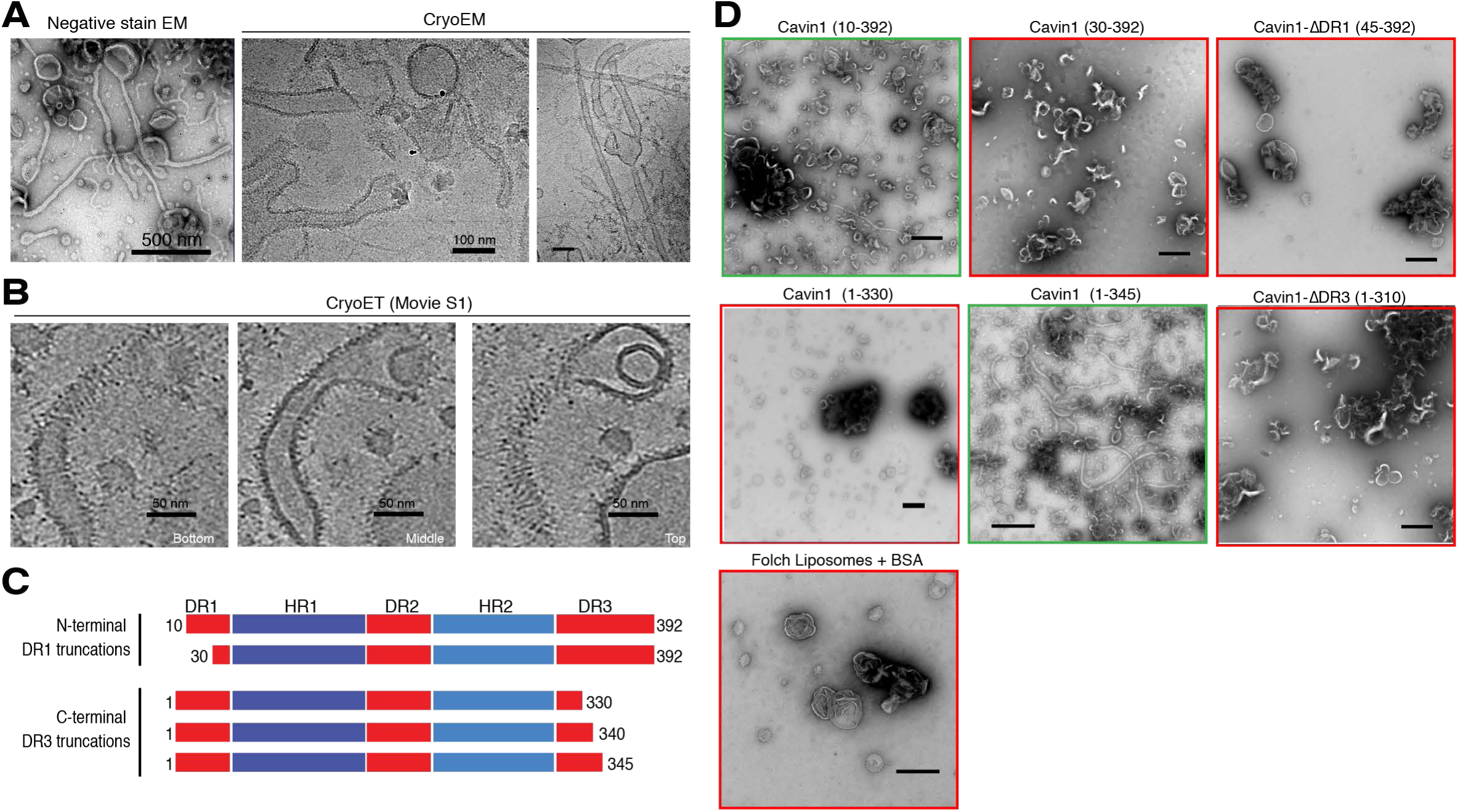
The Cavin1 DR domains are required for membrane remodelling *in vitro.* **(A)** Purified Ub-tagged full length Cavin1 was mixed with Folch 400 nm unilamellar liposomes and analysed by both negative stain EM (1% uranyl acetate) and cryoEM. (**B)** Cryoelectron tomography (CryoET) of Cavin1-coated membrane tubules showing bottom, middle and top sections of three-dimensional projections. Striated protein densities are observed coating the relatively heterogeneous membrane tubules. The full tomogram is shown in **Movie S1**. (**C**) Schematic diagram of Cavin1 and different truncation constructs examined for their ability to remodel membranes *in vitro*. **(D)** Purified Ub-tagged Cavin1 truncations were mixed with Folch 400 nm unilamellar liposomes and analysed by negative stain EM (1% uranyl acetate). Full membrane tubulation and remodelling activity requires residues 1-30 in DR1, and residues 330-345 in DR3. Scale bar = 500 nm.

The importance of Cavin1 DR domains in self-association and LLPS raised the question as to whether they play a role in its ability to physically remodel membranes. To this end, we used the *in vitro* membrane remodelling assay to investigate their importance in generating membrane curvature. We expressed and purified a range of Cavin1 DR domain truncations with an N-terminal His-ubiquitin (HisUb) tag (**Fig. 4C; Fig. S2**) and used the membrane tubulation assay combined with negative stain EM to analyse their ability to remodel mammalian (Folch) synthetic phospholipid membranes (**Fig. 4D**). Complete removal of either the N-terminal DR1 or C-terminal DR3 domains abolished the ability of Cavin1 to tubulate liposomes *in vitro*. Shorter truncations showed that while the N-terminal DR1 deletion Cavin1(10-392) still formed membrane tubules, these were relatively infrequent and of a smaller diameter (∼10 nm), whereas further deletion of N-terminal DR1 sequences in Cavin1(30-392) prevented the formation of membrane tubules altogether. The C-terminal DR3 deletion mutant Cavin1 (1-345) formed membrane tubules similar to full length Cavin1. However, the deletion of further amino acids from the C-terminus in Cavin1(1-330) completely inhibited membrane tubulation. For those DR truncation mutants that lacked membrane remodelling activity we observed instead a propensity to cause liposome clustering. This likely occurs because these Cavin1 constructs can now bind adjacent phospholipid vesicles via multiple positively charged surfaces of the HR1 and HR2 domains, unrestrained by compensating negatively-charged DR1 and DR3 sequences ^11^. Overall, these studies define a core Cavin1 sequence (10-345) required for Cavin1 to efficiently promote membrane curvature.

We next examined the ability of purified GFP-Cavin1 to modulate Folch lipid giant multilamellar vesicles (GMVs) doped with 0.1 mol% fluorescent Bodipy-TMR PI(4,5)P2 analogue. GFP-Cavin1 showed strong localised clustering at the membrane surface compared to other membrane remodelling proteins (**Fig. 5A**) ^57, 58^, and possessed a remarkable membrane sculpting activity as indicated by the rapid collapse of GMVs over a period of several minutes (**Fig. 5B**). We performed similar experiments with Rhodamine B-PE as a fluorescent marker and observed the same protein clustering and membrane sculpting activity (**Fig. S7C**). In contrast, although both the N- and C-terminal DR deletions of Cavin1 still bound efficiently to GMVs, they did not display any significant membrane sculpting activity **(Fig. 5C, 5D**). With Cavin1-ΔDR1 we also often observed a characteristic accumulation of the protein at the interface between adjoining vesicles leading to the clustering of the GMVs (**Fig. 5E**). Overall, these studies using GMVs and SUVs show that while the DR1 and DR3 domains are dispensable for membrane binding, they have an essential role in the ability of Cavin1 to sculpt the curvature of phospholipid membranes.

**Figure 5.**
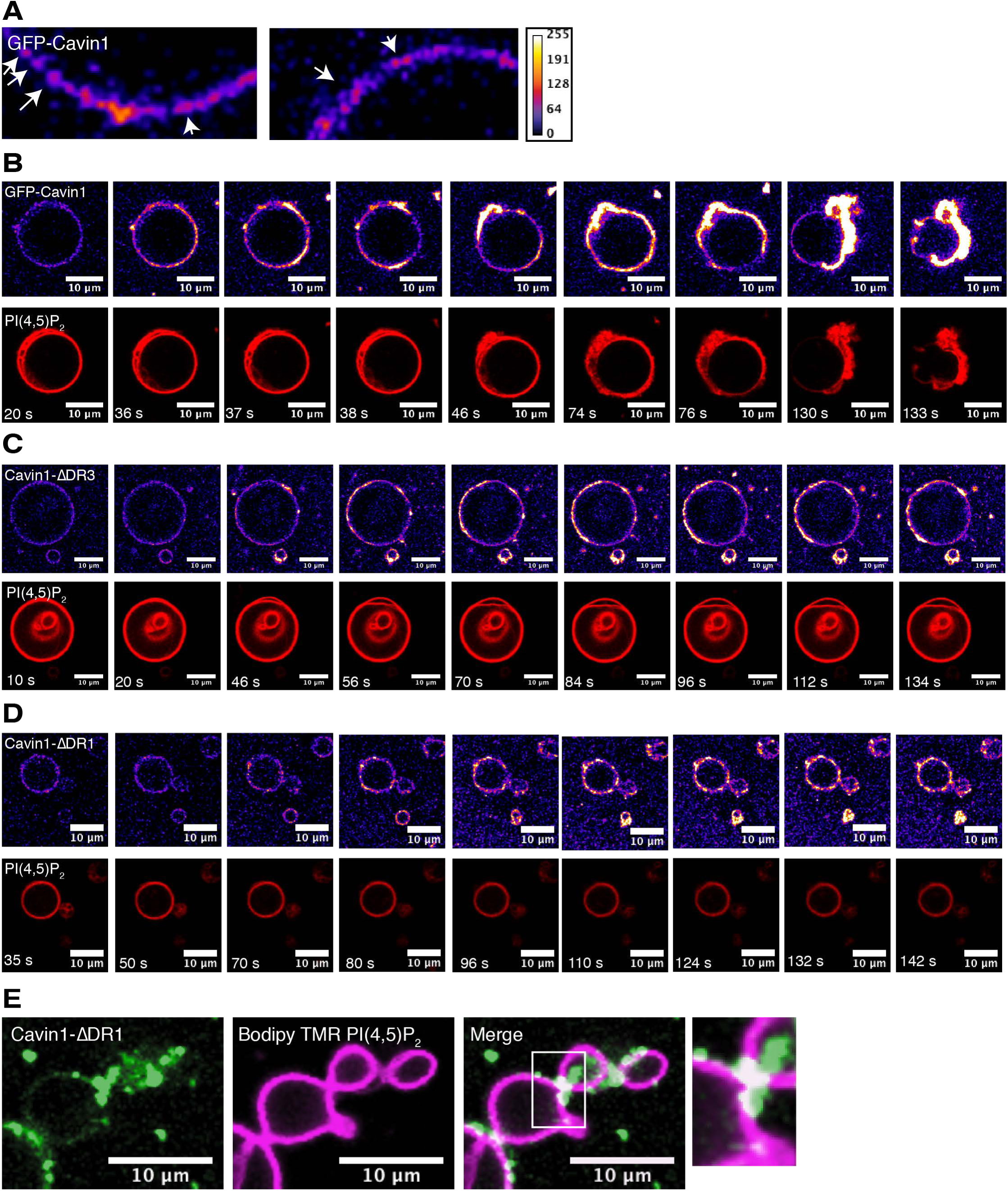
Removing the Cavin1 DR domains prevents deformation of GMV membranes. **(A)** Purified Ub- and GFP-tagged Cavin1 shows strong localised clustering on the surface of Folch giant multilamellar vesicles (GMV) containing Bodipy-TMR-labelled PI(4,5)*P*_2_ (0.1 mol%). Cavin1 **(B)**, Cavin1-ΔDR3 **(C)** or Cavin1-ΔDR1 **(D)** were incubated with Folch GMVs containing Bodipy-TMR-labelled PI(4,5)*P*_2_ (0.1 mol%), allowed to settle on glass coverslips and images were acquired one frame per second. Frame numbers are indicated in PI(4,5)*P*_2_ channel (red). **(E)** GMVs incubated with Cavin1-ΔDR1 were often observed to be tethered to each other with Cavin1-ΔDR1 concentrated at the contact sites. Scale bar = 10 μm.

### Cavin1 disordered sequences are essential for interacting with CAV1 and forming caveolae

Our studies *in vitro* highlight several properties of Cavin1 that are dependent on its disordered sequences. Firstly, DR1 and DR3 of Cavin1 are important for the formation of a large scale associated state and LLPS promoted by electrostatic interactions, and the DR1 domain is required for the dynamic properties of Cavin1 in LLPS; removal of the DR1 domain results in gel formation and prevents its free diffusion within the condensates. The Cavin1-ΔDR1 construct also displays a capacity to bind and cluster membrane vesicles *in vitro*. Secondly, minimal sequences of Cavin1 DR1 and DR3 are required for membrane remodelling. Lastly, the C-terminal DR3 domain of Cavin1 is required for the association with CAV1 in co-mixed liquid droplets *in vitro*.

To examine the importance of Cavin1 disordered N- and C-terminal domains to functional caveola formation, we next analysed the localisation of the DR1 and DR3 truncation mutants in cells using either standard confocal microscopy (**Fig. S8A**) or confocal fluorescence with Airyscan super-resolution imaging (**Fig. 6A**). The prostate cancer PC3 cell line was used, which expresses CAV1 but does not express any members of the Cavin family so that CAV1 is diffusely localised at the plasma membrane ^7, 9^ (**Fig. S8A**). Expression of full-length GFP-Cavin1 in PC3 cells fully restores the formation of caveolae with CAV1 (in the absence of other cavins), providing a functional readout for Cavin1 activity ^7, 9, 10, 59^. Full length GFP-Cavin1 showed a characteristic punctate distribution and co-localised with CAV1 at the plasma membrane (**Fig. S6A; Fig. S8A**). In contrast, after removal of the C-terminal domain GFP-Cavin1-ΔDR3 is unable to promote caveola formation, does not co-localise with CAV1, and now associates extensively with microtubules. This is consistent with a previous report of a similar C-terminal truncated Cavin1 (residues 1-322) in CHO cells ^60^. FRAP analysis of GFP-Cavin1-ΔDR3 on microtubules showed a fast fluorescence recovery, indicating a dynamic exchange with the cytoplasm or diffusion along the microtubules (**Fig 6B, Movie S2**). Interestingly, when microtubules were depolymerised with nocodazole this resulted in redistribution and condensation of GFP-Cavin1-ΔDR3 to form spherical droplets in the cytosol (**Fig. 6C; Fig. S8B; Movie S3**). These also showed fast exchange of protein molecules with the bulk cytoplasm suggestive of liquid-droplet behaviour, and consistent with droplet formation by Cavin1-ΔDR3 *in vitro*. This indicates a dynamic equilibrium exists between cytosolic, liquid droplet and microtubule-associated states of the GFP-Cavin1-ΔDR3 truncation.

**Figure 6.**
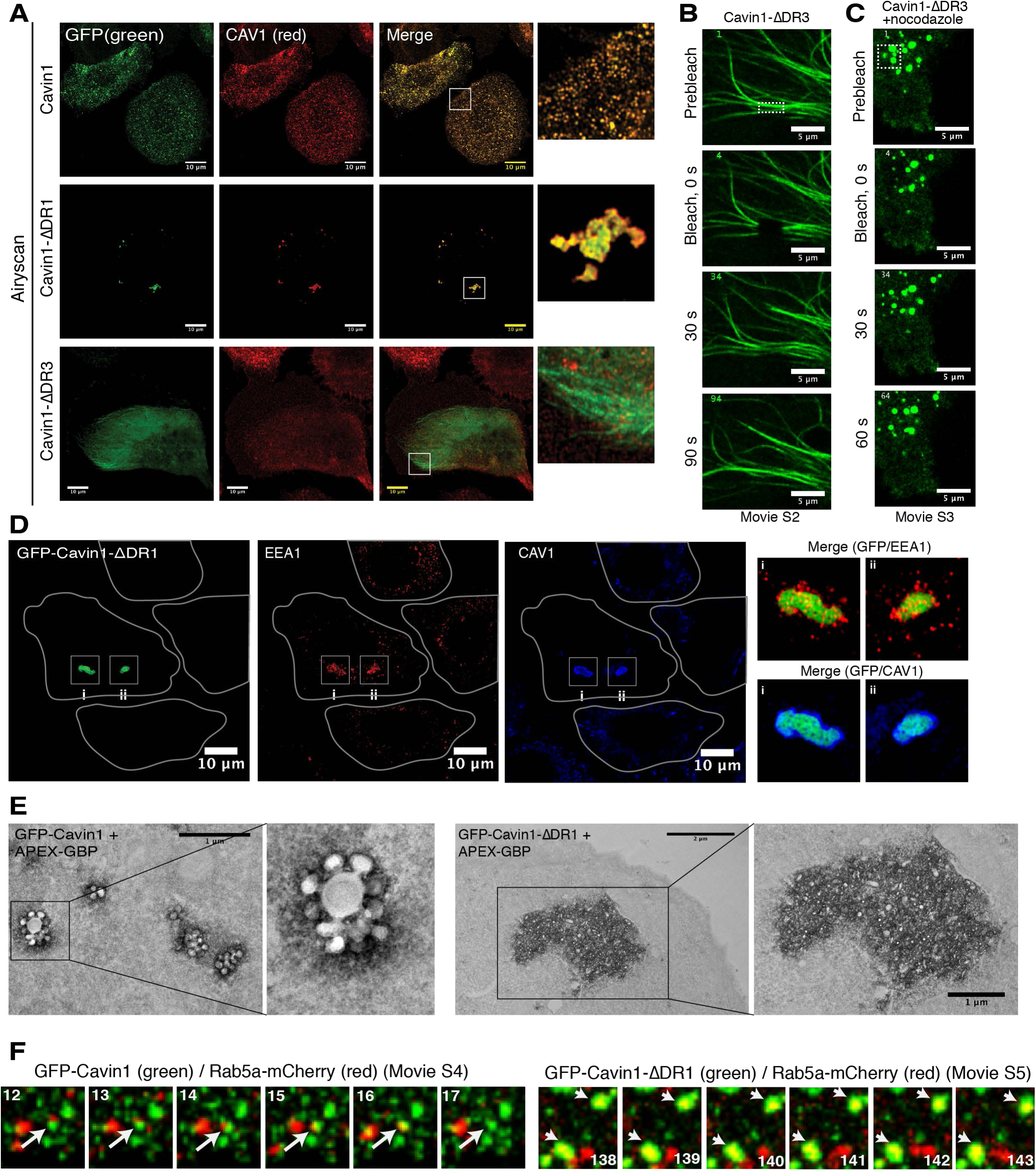
The Cavin1 DR domains are essential for caveola formation with CAV1. (**A**) GFP-tagged Cavin1 and truncations (green) were expressed for 24 h in PC3 cells, fixed and immunolabelled for Caveolin1 (CAV1) (red). Full length Cavin1 forms typical caveola puncta, colocalising with CAV1 at the cell surface. Cavin1-ΔDR1 mutant expression leads to formation of tethered intracellular CAV1-positive clusters. Cavin1-ΔDR3 shows cytoplasmic and microtubule localisation. Images were collected using a Zeiss fast Airyscan microscope. Scale bar = 10 μm. Fluorescence recovery after photobleaching (FRAP) analysis of GFP-Cavin1-ΔDR3 before **(B)** and after **(C)** nocodazole (10 µM) addition. Scale bar – 5 μm **(D)** In PC3 cells GFP-tagged Cavin1-ΔDR1 truncation shows colocalization with the early endosomal marker (EEA1) (red) and CAV1 (blue). Inset shows merge images of GFP-Cavin1-ΔDR1/EEA1 and GFP-Cavin1-ΔDR1/CAV1. Scale bar = 10 μm. (**E**) GFP-tagged Cavin1 and Cavin1-ΔDR1 were visualised in PC3 cells by electron microscopy and labelling of GFP tagged proteins using APEX-GBP staining. Scale bar = 1 μm. **(F)** Live imaging of PC3 cells expressing Rab5a-mCherry with either GFP-Cavin1 or GFP-Cavin1-ΔDR1. Images were acquired one frame per four seconds and frame numbers are indicated in boxes. Arrows indicate mCherry/GFP signal co-localisation or separation event.

Strikingly, expression of GFP-Cavin1-ΔDR1 resulted in the formation of large intracellular structures that also contained endogenous CAV1 (**Fig. 6A; Fig. S8A**). A C-terminal tagged Cavin1-ΔDR1-GFP construct showed similar clusters co-localised with CAV1, confirming this phenotype is not influenced by the location of the GFP tag (**Fig. S8C**). To analyse these structures in more detail, we performed co-localisation experiments of GFP-Cavin1-ΔDR1 with various cellular markers. While no overlap was seen with the Golgi complex, lysosomal or recycling endosomal membrane markers, a significant proportion of GFP-Cavin1-ΔDR1 and endogenous CAV1 were found to colocalise with the early endosomal marker EEA1 (**Fig. S9A**). Airyscan microscopy revealed that clusters of EEA1-positive endosomes surrounded the GFP-Cavin1-ΔDR1 and CAV1-positive structures (**Fig. 6D; Fig. S9B**). We then performed transferrin uptake assays in PC3 cells using transferrin labelled with Alex-488 fluorescent dye. Transferrin positive endosomes showed little overlap with full-length mCherry-Cavin1-positive spots on the cell surface (**Fig. S10**). However, the mCherry-Cavin1-ΔDR1 construct formed intracellular clusters with transferrin positive endosomes surrounding them similar to EEA1. These large intracellular structures were visualised by APEX labelling and electron microscopy imaging ^61^ of GFP-Cavin1-ΔDR1 in PC3 cells, revealing intracellular assemblies consisting of large clusters of vesicles with a surrounding halo of GFP-Cavin1-ΔDR1 labelling (**Fig. 6E**). In contrast, GFP-Cavin1 expression resulted in formation of the characteristic single caveolae and rosettes of caveolae at the plasma membrane as expected (**Fig. 6E**).

We lastly performed live imaging of PC3 cells expressing either GFP-Cavin1 or GFP-Cavin1-ΔDR1 with Rab5a-mCherry as a marker of early endosomes. Caveolae are consistently localised to the trailing edge of migrating cells, where constant membrane remodelling events are occurring ^62^. In migrating PC3 cells we observe dynamic GFP-Cavin1 positive caveola puncta undergoing transient fission and fusion events and kiss-and-run interactions with Rab5a-mCherry positive endosomes similar to previous observations ^63^ (**Fig. 6F**, **Movie S4**). In contrast, GFP-Cavin1-ΔDR1 initially showed plasma membrane puncta fusion events similar to GFP-Cavin1 (imaged at an early 12 h time point following transfection before larger immobile condensates are formed), but over time resulted in formation of the larger structures that stably associated with Rab5a positive endosomes (**Fig. 6F**, **Movie S5**). This suggests that the DR1 domain is important for the dynamics of intracellular trafficking and recycling of caveolae at endosomes. Overall, our results show that disordered sequences of Cavin1 are essential for generating caveolae, but that each DR domain has a distinct function. Removing the C-terminal DR3 domain prevents interaction with CAV1 and results in mis-localisation to the cytoplasm and abnormal association with microtubules. Removing the N-terminal DR1, which results in gel formation and membrane clustering *in vitro,* allows initial caveola formation with CAV1 at the plasma membrane, but then causes subsequent accumulation of aberrant intracellular protein and membrane assemblies with a subset of early endosomes unable to recycle to the plasma membrane.

### Minimal Cavin1 DR sequences needed for membrane remodelling are also essential for caveola formation

Using the series of truncations tested *in vitro* for membrane remodelling activity, we next asked if the same minimal sequences are sufficient for caveola formation in cells. GFP-Cavin1(10-392) showed a relatively normal localisation with CAV1 puncta at the cell surface. However, GFP-Cavin1(30-392) formed large intracellular puncta and clusters that co-localised with CAV1 (**Fig. 7A),** and also showed a partial co-localisation with EEA1 (**Fig. S9C**), similar to Cavin1 with the complete DR1 domain removed. Thus, deletion of the N-terminal DR1 sequence of Cavin1 has a progressive effect on the re-distribution of caveolae from the plasma membrane to intracellular endocytic compartments.

**Figure 7.**
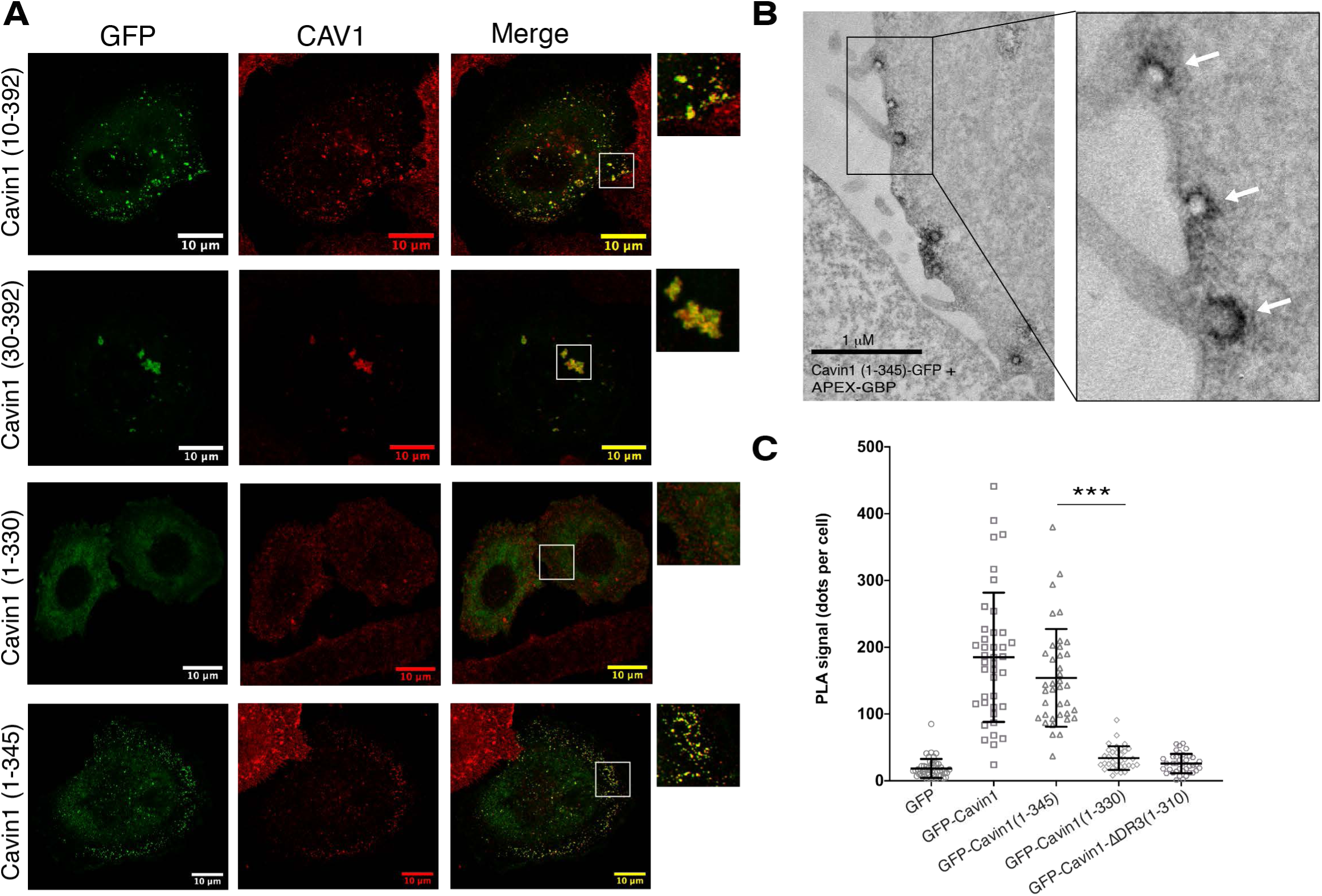
Definition of the minimal DR sequences required for Cavin1 function. **(A)** GFP-tagged Cavin1 DR domain truncation mutants (green) (Fig. 4C) were expressed in PC3 cells and immunolabelled with CAV1 (red). Residues 1-30 in DR1 and 330-345 are required for caveola formation. Scale bar = 10 μm. (**B**) APEX-GBP labelling of GFP tagged Cavin1(1-345) shows normal bulb-shaped caveolae at the plasma membrane. **(C)** Proximity ligation assay (PLA) analyses show that truncation of Cavin1 from the C-terminus beyond residue 345 results in loss of association with CAV1. PLA signal was quantified as dots per cell for specific interaction between GFP-tagged proteins and CAV1, N = 2 (independent biological replicates), n = 10-15 (cells per replicates), Error bars indicate mean ± SD, *** P<0.001.

The C-terminal DR3 truncation GFP-Cavin1(1-345) retained a normal ability to generate plasma membrane puncta that co-localised with CAV1 (**Fig. 7A**), and APEX labelling and electron microscopy of GFP-Cavin1(1-345) showed its typical localisation to caveolae at the plasma membrane (**Fig. 7B**). Further deletion of C-terminal DR3 sequences in GFP-Cavin1(1-330), however, resulted in a total cytosolic redistribution. The C-terminal truncations show that amino acids (346-392) are dispensable for generating caveolae in PC3 cells, while residues 330-345 are essential. Finally, we used a proximity ligation assay (PLA) ^10^ to assess the interactions of Cavin1 C-terminal truncations with CAV1 at the plasma membrane. PLA analyses correlated with the cellular imaging of the GFP constructs, showing that the mutant Cavin1(1-345) can interact with (or is at least in close proximity to) CAV1, while the shorter truncations Cavin1(1-330) and Cavin1(1-310) do not (**Fig. 7C; Fig. S11)**.

### Specific DR sequences are essential for the ability of Cavin1 to form caveolae

The disordered sequences 1-30 and 310-345 in DR1 and DR3 are required for Cavin1 to efficiently self-associate, remodel synthetic phospholipid membranes *in vitro*, and promote caveola formation with CAV1 in cells. To examine these sequences in more detail we generated a series of specific mutations in the DR1, DR2 and DR3 domains in the context of the minimal functional construct Cavin1(1-345) (**Fig. 8A**). Beginning with DR1 (residues 1-30), we first tested whether the acidic amino acids were important by mutating the Glu/Asp residues to alanine (DR1mut1). When GFP-tagged Cavin1(1-345) DR1mut1 was expressed in PC3 cells it formed large intracellular puncta that co-localised with CAV1 (**Fig. 8B**), and also colocalised with a sub-population of EEA1-positive endosomes, but not LAMP1 or GM130 **(Fig. S12A**). APEX labelling and imaging by EM showed clusters of GFP-Cavin1(1-345) DR1mut1 that appeared identical to those formed by either GFP-Cavin1-ΔDR1 or GFP-Cavin1(30-392) (**Fig. S12B**). By FCS, this variant showed a significant increase in diffusivity with respect to wild-type Cavin1(1-345), indicating that its net negative charge is important for self-association (**Fig. S12C**). More precise mutation of Asp/Glu residues in the first ten amino acids of DR1 (DR1mut2) had no qualitative effect on the ability of GFP-Cavin1(1-345) to form caveolae, while altering the Asp/Glu residues in amino acids 10-30 of DR1 (DR1mut3) resulted in the same phenotype as mutant DR1mut1 (or complete deletion of DR1), forming large intracellular clusters with CAV1 (**Fig. 8A and 8B**). We next substituted DR1(1-30) with random Gly/Ser sequences, while maintaining the relative positions of acidic Asp/Glu residues and prolines (DR1mut4). The objective was to determine if any other sequences apart from the acidic side chains contributed to the activity of the domain. In MCF7 cell lysates the DR1mut4 mutant did not show a major difference in diffusivity by FCS compared to wild-type Cavin1(1-345) indicating that only the acidic side-chains in the DR1 region are necessary for self-association (**Fig. S12C**). The subcellular localisation of GFP-Cavin1(1-345) mutant DR1mut4 in PC3 cells also showed co-localisation with CAV1 at the plasma membrane (**Fig. 8B**), indicating that it is the electrostatic properties of the DR1 sequence that are most important for its function and not the specific sequence itself. However, the spacing of acidic residues in DR1 is critical, as complete removal of surrounding sequences (DR1mut5) also results in GFP-Cavin1(1-345) mis-localization. An analogous result was observed for the central DR2 domain of Cavin1, where mutation of the acidic residues (DR2mut6) abolished caveola recruitment in PC3 cells and prevented self-association in MCF7 cell lysates, but altering the surrounding sequences while maintaining negative charges had no effect on caveola formation (DR2mut7) (**Fig. 8C**; **Fig. S12C**). Thus, the presence and the spacing of acidic sequences in DR1 and DR2 are essential for normal caveola formation, but their specific surrounding sequences are not.

**Figure 8.**
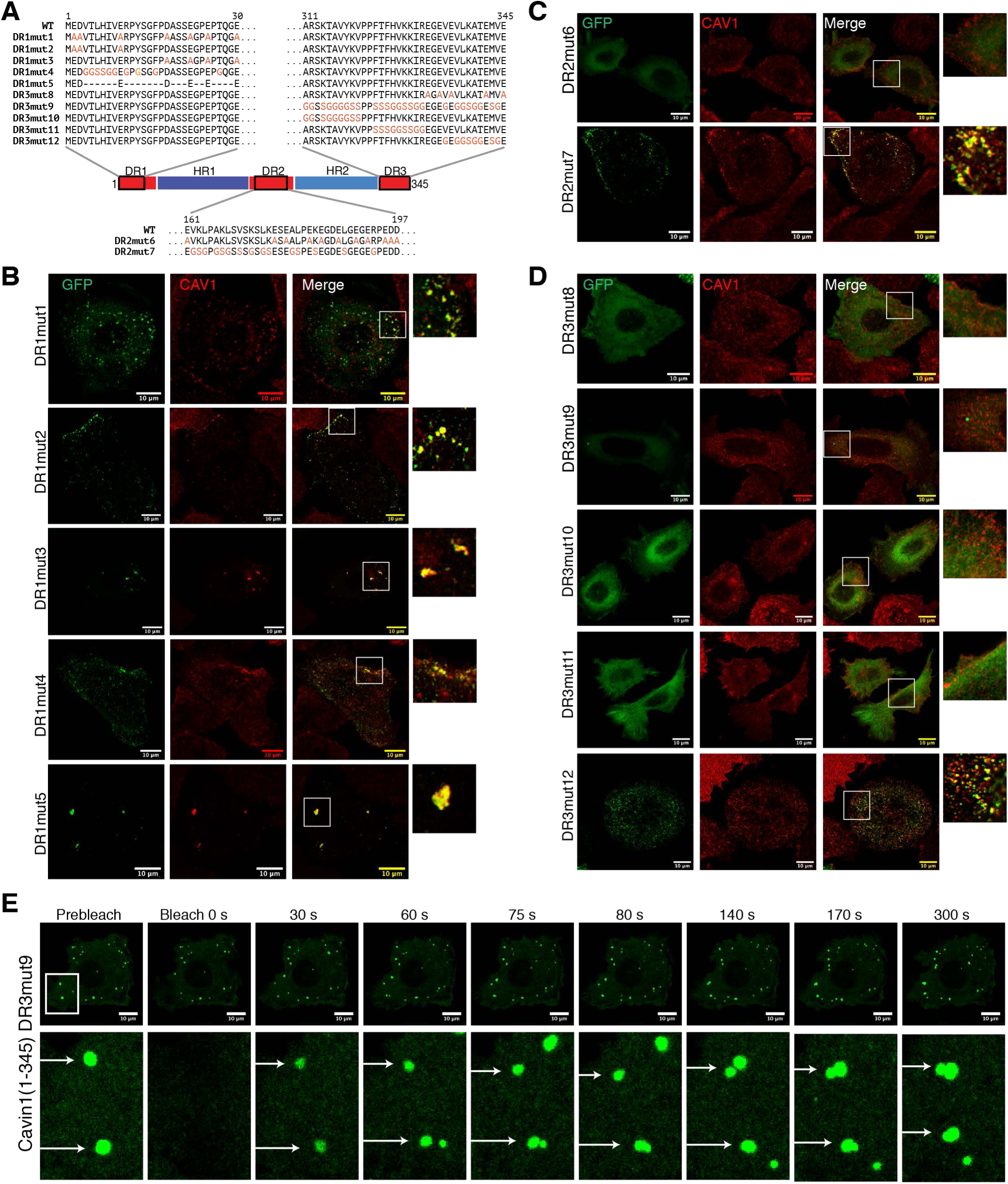
Sequence requirements of the Cavin1 DR domains in caveola assembly. **(A)** Schematic diagram of Cavin1(1-345) with the sequences of the various point mutants indicated. DR1, DR2 and DR3 mutations occur in the regions 1-30, 161-197 and 311-345 respectively. **(B)** GFP-tagged Cavin1(1-345) DR1 domain mutants (green), (**C**) DR2 domain mutants and (**D**) DR3 domain mutants expressed in PC3 cells and immunolabelled with endogenous CAV1 (red). Images in (B), (C) and (D) were by Airyscan confocal microscopy. Scale bar = 10 µm. **(E)** FRAP analysis of Cavin1 (1-345) DR3mut9 mutant showing fast recovery of fluorescence in cytosolic droplets and also droplet fusion events (marked by arrow).

Lastly, we assessed the roles of specific sequences in the essential DR3 region (residues 311-345). The mutation of acidic Asp/Glu residues in GFP-tagged Cavin1(1-345) (DR3mut8) resulted a diffuse cytosolic localisation in PC3 cells (**Fig. 8D**) and prevented self-association in FCS measurements (**Fig. S12C**). These results show that the acidic Glu/Asp residues in the Cavin1 DR1, DR2 and DR3 domains are all essential for oligomeric interactions and forming caveolae with CAV1 at the cell surface. The acidic side-chain mutations result in identical phenotypes to the complete truncation of the DR1 and DR3 domains. In contrast to the DR1 and DR2 domains however, we found that altering everything in DR3 other than Asp/Glu residues (DR3mut9) resulted in a protein with a normal ability to self-associate (**Fig. S12C**), but that was unable to restore caveola formation with CAV1 in PC3 cells (**Fig. 8D**). This protein was generally cytosolic, but in some cells we observed the formation of numerous spherical cytoplasmic structures, that dynamically exchange with the cytosol as shown by FRAP analysis and regularly undergo fusion, suggesting the protein has undergone LLPS and droplet formation (**Fig. 8E; Fig. S12D; Movie S6**). Remarkably however, unlike the complete DR3 deletion purified Cavin1(1-345) DR3mut9 is still able to remodel and tubulate synthetic liposomes *in vitro* (**Fig. S12E**). This shows that while specific sequences in the Cavin1 region 310-345 are dispensable for large scale oligomer formation, LLPS and membrane remodelling, they are still essential for recruitment to caveolae with CAV1 in cells. The acidic side chains in this region, however, are required for all of these functional Cavin1 activities (mutant DR3mut8). To refine this further, we designed three shorter variants of the DR3mut9 mutation, DR3mut10 (311-320), DR3mut11 (321-331), and DR3mut12 (332-345). While mutant DR3mut12 behaved like wild-type Cavin1 and formed normal caveolae, both mutant DR3mut10 and DR3mut11 showed a cytosolic distribution similar to DR3mut9 (**Fig. 8D**). Therefore, specific sequences in the Cavin1 DR3 region 311-331 are specifically required for CAV1 association and caveola formation, while acidic residues within DR3 region (332-345) are essential for promoting electrostatic oligomeric Cavin1 assembly. Lastly, we quantified the co-localization of those Cavin1 DR mutants that still retained prominent association with CAV1 (**Fig S12F**). While the sequences altered in these constructs are not strictly required for caveola formation (e.g. DR1mut2 or DR3mut12) or CAV1 interaction (e.g. DR1mut1, DR1mut3 or DR1mut5), they all showed a marginal reduction in co-localization suggesting they make a minor contribution to Cavin1-CAV1 interactions. Altogether, these studies demonstrate the critical importance of acidic residues in all three DR domains for promoting electrostatic intermolecular interactions and caveola formation; while specific sequences in Cavin1 DR3 region (311-331) are necessary for Cavin1 and CAV1 association for caveola recruitment.

## Discussion

Despite the fact that intrinsically disordered sequences are a prominent and highly conserved feature of all cavins, no previous studies have explicitly addressed their functional importance. We now show that they are essential for caveola formation. In addition, they also regulate the ability of Cavin1 to self-associate and undergo LLPS *in vitro*, where Cavin1 shows the classical properties of LLPS as demonstrated by phase separation that is sensitive to protein concentration, ionic strength, molecular crowding agents, and by the rapid exchange of protein in Cavin1 droplets as shown by FRAP. The sensitivity of this LLPS to salt concentration indicates an electrostatically driven Cavin1 condensation. We demonstrate the distinct roles of the disordered DR domains of Cavin1 in LLPS behaviour, including a mutant protein lacking the DR1 domain that still self-associates but no longer shows the dynamic exchange properties of the full-length protein. In addition, CAV1 was also able to associate with Cavin1 generated liquid droplets, an interaction that is dependent on their mutual disordered sequences. Our cellular studies show that acidic residues in all three Cavin1 disordered sequences (DR1, DR2 and DR3) are essential for generating caveolae with CAV1 at the plasma membrane. Deletion or mutation of these regions in Cavin1 result in mis-localisation and an inability to form plasma membrane caveola invaginations. Interestingly the N- and C-terminal sequences play divergent roles in this process. Deletion of the N-terminal DR1 domain affects caveola dynamics and leads to the formation of large intracellular clusters of Cavin1, CAV1, and endosomal membrane vesicles. In contrast, deletion of the DR3 domain prevents CAV1 association *in vitro* and *in vivo* and results in dynamic microtubule association or cytoplasmic droplet formation. We speculate that Cavin1-ΔDR3 association with microtubules may share mechanistic similarities with the condensation of Tau on microtubules ^64, 65^, or interactions of multivalent positively-charged peptides with the C-terminal acidic tails of tubulin subunits ^66^, but this will require further study.

To better appreciate and visualise the role of the disordered DR domains in Cavin1 activity, we constructed a theoretical structural model of the protein, building on the assumption that the fundamental Cavin1 unit is a homotrimer based on the coiled-coil structure of its N-terminal HR1 domain ^11^ (see Materials and Methods) (**Fig. 9A**). This model points to several interesting features of the Cavin1 protein. Firstly, the combined DR1, DR2 and DR3 domains account for more than 50% of the total Cavin1 sequence. In other words, Cavin1 is not a typical globular protein but rather consists of large random-coil elements tethered by *α*-helical structural cores. Secondly, as suggested by sequence analyses (**Fig. S1**), there is a distinctive electrostatic pattern to the structure, with the *α*- helical domains providing positively charged surfaces for membrane association, and the disordered regions having a generally negatively charged nature. A likely consequence of this is that electrostatic repulsion will cause these DR domains to orient outwards when Cavin1 is in contact with membranes, and we propose they will also form transient electrostatic interactions with the HR domains of neighbouring Cavin1 molecules (**Fig. 9B**). Notably, multiple theoretical and experimental studies have shown that the sequence-specific electrostatically driven interactions between disordered proteins can lead to LLPS and high affinity protein complex formation under physiological conditions, with the tendency to phase separate (or undergo ‘complex coacervation’) increasing as the ‘blockiness’ of the charge distribution increases ^67–72^.

**Figure 9.**
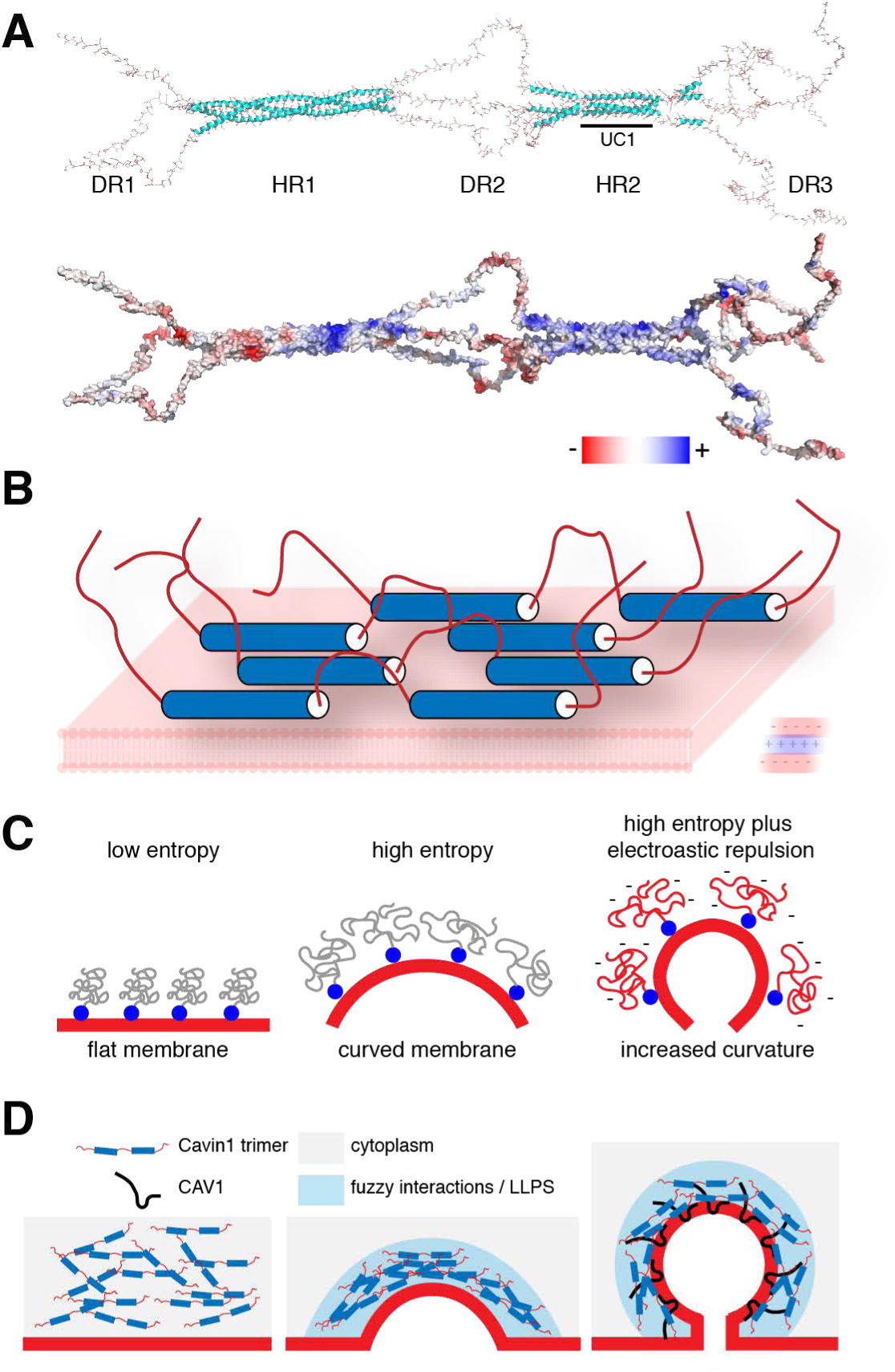
Model for the role of Cavin1 DR domains in LLPS and caveola formation. **(A)** Structural model of a Cavin1 homotrimeric assembly. The trimeric HR1 coiled-coil domain is derived from the crystal structure of the mouse Cavin1 HR1 domain ^11^, the UC1 and HR2 domains are modelled as described previously ^10^, and the DR domains are modelled as random coil structures (see Methods for further details). The structure is shown in ribbon diagram (top) and with an electrostatic surface representation (bottom). **(B)** Proposed orientation of Cavin1 proteins on the membrane surface, with membrane-binding HR1 and HR2 domains associated with the phospholipid bilayer and negatively charged DR sequences directed outwards due to electrostatic repulsion. (**C**) Potential role of Cavin1 disordered sequences in membrane curvature generation due to steric crowding. This concept is largely derived from previous studies of other membrane-associated proteins ^73, 74^. (**D**) Potential role of Cavin1 fuzzy interactions and LLPS in membrane curvature generation, CAV1 interaction and caveola formation.

Our studies of the DR domains of Cavin1 confirm that the acidic residues within these domains are essential for the formation of caveolae in cells and promotion of membrane remodelling *in vitro*. By what mechanism might the DR domains contribute to these membrane sculpting activities? Several recent studies have demonstrated the ability of intrinsically disordered sequences to generate membrane curvature when coupled to membrane binding domains ^57, 73–75^. This is caused by molecular crowding of the disordered sequences leading the proteins to partition with curved or convex membranes so as to increase their conformational entropy; and this can also be enhanced by electrostatic repulsive forces both between the disordered domains and with the membrane itself (**Fig. 9C**). One possible mechanism we can propose for Cavin1-driven membrane curvature is that negatively charged DR sequences and positively charged HR regions of Cavin1 combine to promote self-association, membrane interaction and protein crowding at the membrane surface leading to subsequent membrane bending. In the absence of CAV1 and at high protein concentrations *in vitro*, or under certain conditions in cells, Cavin1 can generate arrays of protein oligomers to form membrane tubules. Under normal conditions however, the process of generating membrane curvature is tightly regulated by CAV1, EHD2 and Pacsin2, and also specific membrane lipids, to restrict Cavin1 remodelling activity only to caveolae. We see an almost complete correlation between the ability of different Cavin1 truncations and mutants to tubulate membranes *in vitro* and the ability to form caveolae *in situ*. The notable exception to this is that alteration of sequences in the DR3 region 310-331 does not affect the ability of Cavin1 to assemble into oligomers and efficiently tubulate synthetic membranes, but still results in a failure to generate caveolae in cells. This implies these specific sequences in the Cavin1 DR3 region are additionally required for Cavin1 recruitment to CAV1-positive membrane domains through interactions with the disordered CAV1 N-terminus.

A second mechanism for membrane curvature suggested by our results (and not mutually exclusive with a role for molecular crowding) is the formation of phase-separated Cavin1 domains that incorporate membrane-embedded CAV1. Intrinsically disordered regions of proteins have gained significant attention for their ability to promote LLPS, or biomolecular condensation, with important biological functions such as stress granule formation, assembly of nuclear sub-structures and sensing changes in cellular homeostasis ^19, 76^. The plasma membrane and surfaces of intracellular compartments including the ER and lysosomes have been found to play a role in LLPS, acting as sites of droplet nucleation or as platforms for transport of phase-separated assemblies for example ^23–34^. It has also recently been proposed that biomolecular condensates associated with phospholipid membranes might possess emergent mechanical properties that can result in membrane curvature generation ^35–37^. This is depicted in schematic form in **Fig. 9D**. Here we have shown for the first time that purified Cavin1 can readily undergo LLPS under near physiological conditions and is able to recruit CAV1 through interactions involving LLPS. The DR1 and DR3 domains contribute to this process, although neither domain is strictly essential. Indeed, mutations in DR3 that maintain its negative charge but prevent CAV1 interaction at the plasma membrane actually promote GFP-Cavin1 liquid droplet formation in cells. Notably, removal of the DR1 domain results in apparent gel formation rather than liquid droplet assembly *in vitro*, and within cells results in a striking accumulation of large intracellular structures that also contain CAV1. These are formed by endocytic redistribution of caveola structures from the cell surface and accumulation with early endosomal membranes. Caveolae, positive for both CAV1 and Cavin1, have been shown to bud from the plasma membrane and fuse with early endosomal compartments ^46, 77–81^, and this would almost certainly require dynamic remodelling of the protein coat to allow the fusion process to occur. We postulate that the intracellular structures we observe with Cavin1-ΔDR1 are formed by internalised caveolae, which have become trapped during the stage of early endosomal fusion. This may be due to the DR1-truncated Cavin1 being unable to undergo normal dynamic exchange, as suggested by its gel-forming properties and its propensity to cluster membrane vesicles, causing inhibition or slowing of the docking and fusion with the early endosome in a process involving EEA1 and Rab5a ^80, 82^.

Our data indicates that the assembly of caveolae by Cavin1 strictly depends on a ‘fuzzy’ network of interactions promoted by electrostatic associations, with an essential role for the intrinsically disordered DR domains of Cavin1 in self-association, CAV1 interaction, membrane remodelling and ultimately caveola formation. Fuzzy interactions are defined broadly as those that involve dynamic, exchanging, multivalent interactions with varying degrees of protein disorder or structural ambiguity ^83–85^. This provides versatility and reversibility in protein-protein interactions, and such fuzzy interactions are also proposed to be a driver of protein phase transitions ^19^. One of the historically consistent observations regarding caveolae is that they do not possess an obvious or highly ordered coat morphology akin to clathrin or COP-coated vesicles. In previous studies of caveola architecture it is notable that while some recurring structures are observed, the general appearance of the caveola surface is extremely heterogeneous ^54–56, 86^. Our model for caveola assembly and structure differs markedly from other classical membrane coats such as clathrin or COPI and COPII, which are built from symmetrical arrays of structured protein domain interactions. While structural elements of cavins and caveolins will likely produce semi-regular spacings between the building blocks, the flexible nature of the disordered domains that provide the ‘glue’ for caveola assembly mean that the overall organisation of the coat will be highly dynamic. Caveola formation is the result of multiple low affinity fuzzy interactions between Cavin1, CAV1 and membrane lipids, and we propose that this leads to a metastability in caveola structure that is important in both the dynamic cycling of caveolae through the endocytic pathway and also for their ability to respond to stresses by rapid disassembly.

## Materials and methods

### Cell lines maintenance and materials

PC3 cells were maintained in RPMI medium (Gibco® Life technologies) supplemented with 10% fetal bovine serum (FBS) and Penicillin/Streptomycin. Cell lines were sourced from ATCC and tested fortnightly for mycoplasma contamination. For all experiments, 2 X 10^5^ PC3 or MCF7 cells were plated in either 6 well culture dishes (Nunc™, Cat. No. 140675, Culture area - 9.6 cm^2^) or glass bottom 35 mm dishes (ibidi, No. 1.5 glass coverslip bottom Cat No. 81218) or 35mm tissue culture dishes (TPP**^®^** 93040, culture area - 9.2 cm^2^). Antibodies used were as follows, rabbit polyclonal anti- Caveolin1 (BD Transduction Laboratories, Cat. No. 610060), mouse monoclonal anti-GFP (Roche Diagnostics Cat. No. 11814460001), Donkey anti-Rabbit IgG (H+L) Secondary Antibody Alexa Fluor® 555 conjugate (Thremo Fisher Scientific, Cat No. A31572). Mouse monoclonal anti-tubulin (Anti-alpha Tubulin antibody [DM1A] - Abcam (ab7291)). Folch lipids were obtained from Sigma Aldrich Folch I fraction (B1502).

### Molecular cloning and plasmids

For Recombinant protein expression in *E. coli* two vectors (pHUE and pOPINE-GFP) were used to generate Cavin1 DR domain variants summarised in **Fig. S1**. pHUE vector was used to generate N-terminal 6X-Histidine-Ubiquitin tagged DR domain variants of Cavin1 using overlap extension polymerase chain reaction (OE-PCR) technique at SacII restriction enzyme site ^87^. GFP tagged cavin DR domain variants were generated using pOPINE-GFP vector (in house vector with pOPINE backbone containing GFP) BamHI restriction enzyme site with N-terminal 6X-Histidine-Ubiquitin tag and C-terminal GFP tag using OE-PCR ^88^. For mammalian cell expression constructs, eGFPC1 and eGFPN1 vectors were used to generate respective DR domain Cavin1 mutants summarised in **Fig. S2**. Specific Cavin1 (1-345) DR domain genes (summarised in **Fig. 8A**) and all mCherry tagged CAV1 genes were artificially synthesized (Gene Universal) and selective genes were subsequently cloned into eGFPC1 and pHUE vectors using OE-PCR for mammalian and bacterial expression respectively.

### Recombinant protein expression and purification

Recombinant protein expression was performed using *Eschericia coli* strain Rosetta™ 2 (DE3) (Novagen) (Merck Cat. No 71403). Protein expression was always performed using freshly transformed chemically competent *E. coli* Rosetta 2 cells with respective plasmids. Cell were propogated in either LB or TB media and protein expression was performed by inducing with 0.5 mM IPTG (Isopropyl ß-D-1-thiogalactopyranoside, Bioline, Cat No. BIO-37036) overnight at 18^0^C. Next day, cells were harvested in 20 mM HepesKOH (pH 7.6), 500 mM NaCl (500GF buffer) with addition of benzamidine hydrochloride (Sigma Aldrich, B6506) and cOmplete™, EDTA-free Protease Inhibitor Cocktail Roche (Sigma Aldrich, 4693132001). Cleared cell lysates were prepared using a continuous flow cell disruptor (Constant Systems Limited, UK) at pressure range 25 – 30 kPsi with subsequent addition of 0.5 % w/v Triton X-100 (Cavin1 purification) or n-dodecyl β-D-maltoside (DDM) (1.2 mM) (MBP-GBP-CAV1 purification) and 5 mM imidazole (Sigma Aldrich, 792527) followed by centrifugation 35,000x *g* for 30 min. Purification of 6X-Histidine tagged cavin proteins was done using TALON metal affinity resin (ClonTech, Scientifix Cat No. 635503). Talon resin was thoroughly washed with 500GF buffer containing 5 mM imidazole to remove detergent and non-specifically bound proteins, and elution was performed in 500GF buffer containing 300 mM imidazole. Protein samples were immediately loaded on size exclusion chromatography column Superose 6 10/30GL pre-equilibrated with 20 mM HepesKOH pH7.6, 150 mM NaCl (150GF buffer) or 150GF buffer with 1.2mM DDM detergent. Size exclusion profiles of purified Cavin truncation mutants are shown in **Fig. S11**. The purified protein used in assays (marked with arrows) appears to be slightly truncated but forms part of megadalton size full length protein complex eluting in higher molecular weight fractions (**Fig. S11**). This partial truncation can be due to presence of multiple protease sensitive PEST (proline, glutamate, serine, threonine) regions in DR sequences of Cavin1 ^9^. There has been evidence for the presence of truncated species of Cavin1 bound to native caveolae in cells suggesting that this might be an inherent property of this protein regardless of its source of expression ^7, 89^.

Purification of mammalian Cavin1 was performed by Transfecting GFP-tagged Cavin1 using polyethylenimine (PEI) transfection reagent (Sigma-Aldrich Cat. No. 408727) with 1:4 w/w ratio (DNA:PEI) and cells were harvested 24 h post-transfection. Cell lysis was performed in 20 mM Hepes-KOH pH 7.6, 500 mM NaCl buffer containing 1% Triton X-100 with three times 3-s sonication pulse at output power 10. Lysate was then centrifuged at 5,000X g for 10 min, and supernatant fraction was incubated with purified GFP nanobody tagged with MBP for 30 min at 4°C. Finally, lysate nanobody mixture was incubated with amylose resin (NEB Cat No. E8021L) for 2 h at 4°C. Amylose resin was then extensively washed with 20 mM Hepes-KOH pH 7.6, 500 mM NaCl buffer containing 1% Triton X-100, and elution was performed in 20 mM Hepes-KOH pH 7.6, 500 mM NaCl buffer containing 20 mM Maltose (Sigma-Aldrich Cat. No. M2250). Elute was concentrated with subsequent addition of Prescission protease to remove MBP tag and finally loaded on Superose 6 10/ 30GL size exclusion column equilibrated in 20 mM Hepes-KOH pH 7.6, 150 mM NaCl. Concentration of gel filtration elute was avoided as it leads to precipitation in lower salt content.

### Fluorescence correlation spectroscopy (FCS) analysis

FCS analysis was performed on a Zeiss LSM 710 confocal microscope equipped with 40x/1.2W C-Apo lens and twin BiG GaAsP detectors capable of single molecule detection. Briefly, MCF7 cells were lysed in a buffer (200 μl) containing 25 mM Tris (pH 7.4), 150 mM NaCl while passing through a 27G needle 6 times. Lysates were then centrifuged at 17,000x g for 10 min to remove cell debris. Supernatant was then used for FCS analysis. Purified GFP tagged cavin proteins were prepared for FCS by dilution of respective stock solutions of cavin truncates in either 500GF or 150GF buffer to achieve 0.1 µM protein concentration with subsequent centrifugation at 17,000 X g for 10 min. At the beginning of each FCS session on a Zeiss LSM 710, pinhole calibration was done with BODIPY-FL maleimide dye (Cat. No. B30466). Subsequently, diffusion time for three dyes that differ in molecular weight and particle size BODIPY-FL maleimide (∼24 μs), BODIPY-FL iodoacetamide (∼22 μs) and TAMRA DIBO (∼37 μs) was measured for each session (**Fig. S11I**). FCS measurement for each GFP tagged Cavin protein was then done for 10 s and repeated 10 times with a binning time of 200 ns. FCS measurements showing presence of aggregates were removed from analysis. The autocorrelation function G(τ) was fitted using a predefined isotropic 3D translational diffusion Gd(τ) model from the ConFoCor model tool with fixed amplitude (A) and structural parameter, G(τ) = 1 + A * Gd(τ). The diffusion coefficient or diffusivity (µs^2^Sec^-1^) for each measurement was exported from the Zeiss analysis program and plotted for all constructs in Graph pad Prism software. Hydrodynamic radius calculations were done using Stokes-Einstein equation with basic assumption of perfect spherical object diffusion. Stokes – Einstein equation; D = KBT/6πnr, Where, KB – Boltzmann constant, T – Temperature (298K), π – pi (3.14), n – dynamic viscosity (Pa.S) and r – hydrodynamic (Stokes) radius of spherical particle.

### In vitro phase separation assays

Purified GFP-tagged Cavin1 proteins, or mixtures of mCherry-CAV1 and GFP-Cavin1 proteins, were diluted to respective protein and/or salt concentrations prior to addition of dextran T-500 (Pharmacia). Dextran solution was added on the top of protein solution without any mixing to allow natural diffusion of dextran. Image acquisition and fluorescence recovery after photobleaching (FRAP) assays were performed after 2 min wait period to allow phase separated droplets to settle. Phase separation analysis was done within 10 min post addition of dextran. Non-bleaching image acquisition conditions were established before performing FRAP assay. FRAP analysis was done by bleaching rectangular area (2 μm X 1 μm approximately) within protein droplet using 488 nm Argon laser and subsequent image acquisition was done one frame per second. Recovery curves from different proteins were normalised without acquisition bleaching correction using formula [F_(T)_ –F_(postbleachT=0)_]/[F_(Prebleach)_ – F_(postbleachT=0)_]. Normalised data points were used to perform non-linear exponential recovery fit using equations within ImageJ 1.50g or Prism version 8 to obtain half-life value for fluorescence recovery of respective protein.

### Liposome preparation and in vitro membrane tubulation assay

Liposomes were prepared by mixing of 10 µL 10 mM stock solution of Folch lipids (bovine brain extract lipid - Folch I fraction Sigma Aldrich B1502) with 50 µL chloroform in a round-bottom flask. The mixtures were dried gently by a stream of nitrogen first and under vacuum overnight thereafter. Liposomes were rehydrated in 500 µL 150GF buffer followed by repetitive freeze-thaw cycles for 3 – 5 times, using first a mixture of dry ice and acetone followed with 60°C water. The liposomes were then extruded through a 400-nm polycarbonate membrane 21 times using an Avanti mini-extruder to generate large unilamellar lipid vesicles (LUVs).

A 5 µl volume of purified Cavin1 variants [∼ 0.1 mg/ml (1.5 – 2 µM)] was mixed with 5 µl 200 µM liposomes for 1 – 3 min at room temperature. Samples were then quickly spotted onto formvar-carbon coated electron microscopy grids (Cu/Pd grids 200 mesh hexagonal – ProSciTech - GCU-PD200H) for 10 s and excess samples were removed by blotting at corner using Whatman filter paper. This is followed by 2 - 3 distilled water washes in similar fashion and subsequent application of 1% uranyl acetate stain. The excess of stain was removed by blotting and grids were allowed to air dry for a while before viewing under the electron microscope. Final images were acquired on JEOL 1011 electron microscope at 80 kV.

### Giant multilamellar vesicle (GMV) experiments

Giant multilamellar vesicles (GMV) were prepared using electro-formation method described before^90^. Briefly, lipids mixture dissolved in chloroform / methanol solution was gently applied to indium-tin-oxide coated glass slide (Sigma Aldrich Cat. No. 636908) as multiple layers. This solution was then dried under constant stream of nitrogen to remove organic solvent and further dried under vacuum overnight. Next day, electro-formation was performed at 50^0^C in 150GF buffer for 1 hr. Vesicles were used immediately for experiments.

### Cryoelectron microscopy / tomography of Cavin1 coated membrane tubules

Liposome tubulation reaction was assembled as described in the previous section and subjected to vitrification after a 1 – 3 min incubation period. For vitrification, the sample was applied to Lacey carbon grids (EMS, Hatfield, PA,USA) using a Vitrobot Mark II (FEI, Eindhoven, NL) plunge freezer with 4 µl of sample, 6 s blotting time and a -3 mm offset at 24^0^C and 100% humidity. Images were collected on a Tecnai G2 F30 TEM (FEI, Eindhoven, NL) operated at 300 kV at a magnification of 12,000X with 5 µm defocus. Images were recorded on a Gatan K2 summit camera in counting mode for a final pixel size of 3.556 Å per pixel. Images were processed in either IMOD (version 4.9) or ImageJ.

Tilt-series were acquired on a Talos Arctica TEM (Thermo Fisher Scientific-FEI, Eindhoven, NL) operated at 200 kV and at a magnification of 45,000x (final pixel size 3.11 Å per pixel). Images were recorded using the microscope software Tomography (Thermo Fisher Scientific-FEI, NL) on Falcon 3 (Thermo Fisher Scientific-FEI, NL) camera operated in counting mode at an angular range of -60 ° to 60 ° in a bidirectional fashion and at an angular increment of 2°. The defocus was set to -5 µm. Unbinned movies of 8 frames with a set dose rate of ∼1.7 e/Å^2^ were acquired and tomographic reconstructions were generated using the weighted back-projection method in IMOD (https://bio3d.colorado.edu/imod/ version 4.9).

### Electron microscopy processing of PC3 cells

PC3 cells were plated onto 30 mm tissue culture dishes and allowed to adhere to dishes for 48 h prior to transfection. Cells were then co-transfected with APEX-GBP and respective cavin1 mutant constructs. 24 h post transfection, cells were fixed with 2.5% glutaraldehyde in 0.1 M sodium cacodylate buffer (cacodylate) (pH7.4) for 1 h. DAB (3’3-diaminobenzidine tetrahydrochloride, Sigma-Aldrich Cat. No. D5905) reaction was then performed as follows. Briefly, cells were washed with DAB/cacodylate mixture (DAB final concentration – 1 mg/ml) for 2 mins, then treated with DAB/cacodylate + 5.88 mM H2O2 (hydrogen peroxide, Sigma-Aldrich Cat. No. H1009) for 20 mins. Cells were then washed with 0.1 M sodium cacodylate buffer and contrasted with 1% osmium tetroxide for 2 mins. Cells were then embedded in LX112 resin and thin sections were cut as described previously ^61^. Images were acquired on JEOL 1011 electron microscope fitted with a Morada CCD camera (Olympus) under the control of iTEM software and operated at 80kV.

### Immunofluorescence analysis, live cell imaging and Proximity ligation assay (PLA)

PC3 cells were grown at about ∼50% confluency in RPMI 1640 medium supplemented with 10% FBS. Cells were then transfected with respective Cavin1 mutants using Lipofectamine 3000 (Invitrogen) as per manufacturer’s instructions. Cells were fixed 24 h post transfection with 4% paraformaldehyde in phosphate-buffered saline (PBS) at 4°C and subsequently permeabilised with 0.1% Triton X-100 in PBS for 7 mins. Cells were probed with CAV1 antibody (Dilution 1:600) and anti-Rabbit secondary antibody Alexa Fluor® 561 conjugate (Dilution 1:400). For Transferrin uptake assays, PC3 cells expressing either GFP-Cavin1 or GFP-Cavin1-ΔDR1 were incubated with transferrin labelled with Alexa-488 (5 μg/ml) for 1h at 37 °C. Cells were then washed three times with ice cold PBS and cell were subsequently fixed with 4% paraformaldehyde in PBS for all experiments except live imaging. Cholesterol addition experiments were performed in MCF7 cells expressing GFP tagged Cavin1 with serum starvation (Serum free DMEM + 1% BSA, 1h) prior to the addition of water soluble analog of Cholesterol (Sigma-Aldrich Cat. No. C4951). Cells were incubated in DMEM media containing Cholesterol for 40 min at 37°C with immediate fixation using 4% paraformaldehyde in phosphate-buffered saline (PBS) at room temperature. Confocal images (1024X1024) were acquired on Zeiss inverted LSM 880 coupled with fast airyscan detector (Carl Zeiss, Inc) equipped with 63X oil immersion objective, NA 1.4. Images were acquired at different laser power for GFP tagged truncation mutants and detector gain settings in order to avoid oversaturation of pixels. All images were processed for brightness/contrast (histogram) adjustment for visualisation using ImageJ. For live cell imaging and FRAP analysis, cells were plated on glass bottom (No. 1.5) petri dishes (ibidi) and allowed to grown at about ∼70% confluency and transfected with respective Cavin1 mutants. For bleaching, 488 nm laser at 100% attenuation power was used for 20 iterations and subsequent imaging was done at one frame per second. Airyscan processing was done automatically in Zeiss software (ZEN 2.3). For PLA, PC3 cells were processed as described previously ^10^. Images were then acquired on Zeiss LSM 710 and LSM 880 confocal microscope (Carl Zeiss, Inc) equipped with 63X oil immersion objective and quantitation of PLA dots per cell was performed using find maxima function in ImageJ with offset of 25. For quantitative co-localization, images (1024X1024) were acquired on Zeiss inverted LSM 880 in confocal mode and Pearson’s coefficient calculation was done using colo2 macro using imageJ (https://imagej.net/Coloc_2),

### Constructing a structural model of mouse Cavin1

A structural model of mouse Cavin1 was built manually based both on known structures of the mouse Cavin1 HR1 domain ^11^ (PDB ID 4QKV), the previous model of the Cavin1 undecad UC1 region ^10^, and secondary structure prediction of the Cavin1 protein carefully cross-referenced to several Cavin1 homologues and other Cavin family members ^11^. Based on the homotrimeric coiled-coil structure of the HR1 domain we constructed our model under the assumption that a single Cavin1 complex would consist of three separate Cavin1 chains. The secondary structure predictions and previous crystal structure led us to define the following regions of Cavin1 as either *α*-helical or random-coil; DR1, residues 1-48, random-coil; HR1, residues 49-147, *α*-helical (based on PDB 4QKV of mouse Cavin1 HR1); DR2, residues 148-218, random-coil; HR2, *α*-helical for residues 219-242, random-coil for residues 243-244, *α*-helical for residues 245-278 (model from ^10^), random-coil for residues 279-286, *α*-helical for residues 287-297; DR3, residues 298-392, random coil. Stretches of random-coil were built and added to *α*-helical domains manually in COOT Version 0.8.2 ^91^, and the final model was subjected to simple geometry regularisation in PHENIX Version 1.14 ^92^. Structural images and electrostatic surface representations were rendered with PYMOL Version 2.3.1.

### Statistical analyses

Statistical analysis and P value calculations were performed by one-way ANOVA using graph pad Prism software.

## Acknowledgements

This work was supported by grants from the National Health and Medical Research Council of Australia (NHMRC) (to RGP grant number APP569542 and APP1037320) and the Australian Research Council (ARC) (to BMC, grant number DP120101298). RGP is supported by an NHMRC Senior Principal Research Fellowship from the NHMRC (APP1058565) and by the Australian Research Council Centre of Excellence in Convergent Bio-Nano Science and Technology (R.G. Parton). BMC is supported by an NHMRC Senior Research Fellowship (APP1136021). Confocal microscopy was performed at the Australian Cancer Research Foundation (ACRF)/Institute for Molecular Bioscience (IMB) Dynamic Imaging Facility for Cancer Biology, established with funding from the ACRF. The authors acknowledge the use of the Cryo Electron Microscopy Facility through the Victor Chang Innovation Centre, funded by the NSW government, and the Electron Microscope Unit within the Mark Wainwright Analytical Centre at UNSW Sydney.

## Author contributions

BMC and RGP conceived the project, study and acquired funding. VT performed molecular cloning, in vitro protein purification, in vitro, cellular assays, live imaging and FCS experiments. GY assisted in molecular cloning and in vitro protein purification. KAM assisted in cellular and phase separation assay. OK performed preliminary FCS analysis of cavins in the eukaryotic Leishmania terentolae cell free lysate system. NC initiated cholesterol addition experiments completed by VT. JR performed transferrin uptake assay, cellular processing for electron microscopy and electron microscopy image acquisition. NA and MF performed tomography data acquisition, and trained VT in cryo-EM methods. All authors commented on the manuscript. VT, RGP and BMC wrote the manuscript.

## Data availability

Source data for Figs. 1, 2, 6, and 7 are provided in Table S1. The data that support the findings of this study are available from the corresponding author on request.

## Supplementary Information

**Figure S1.**
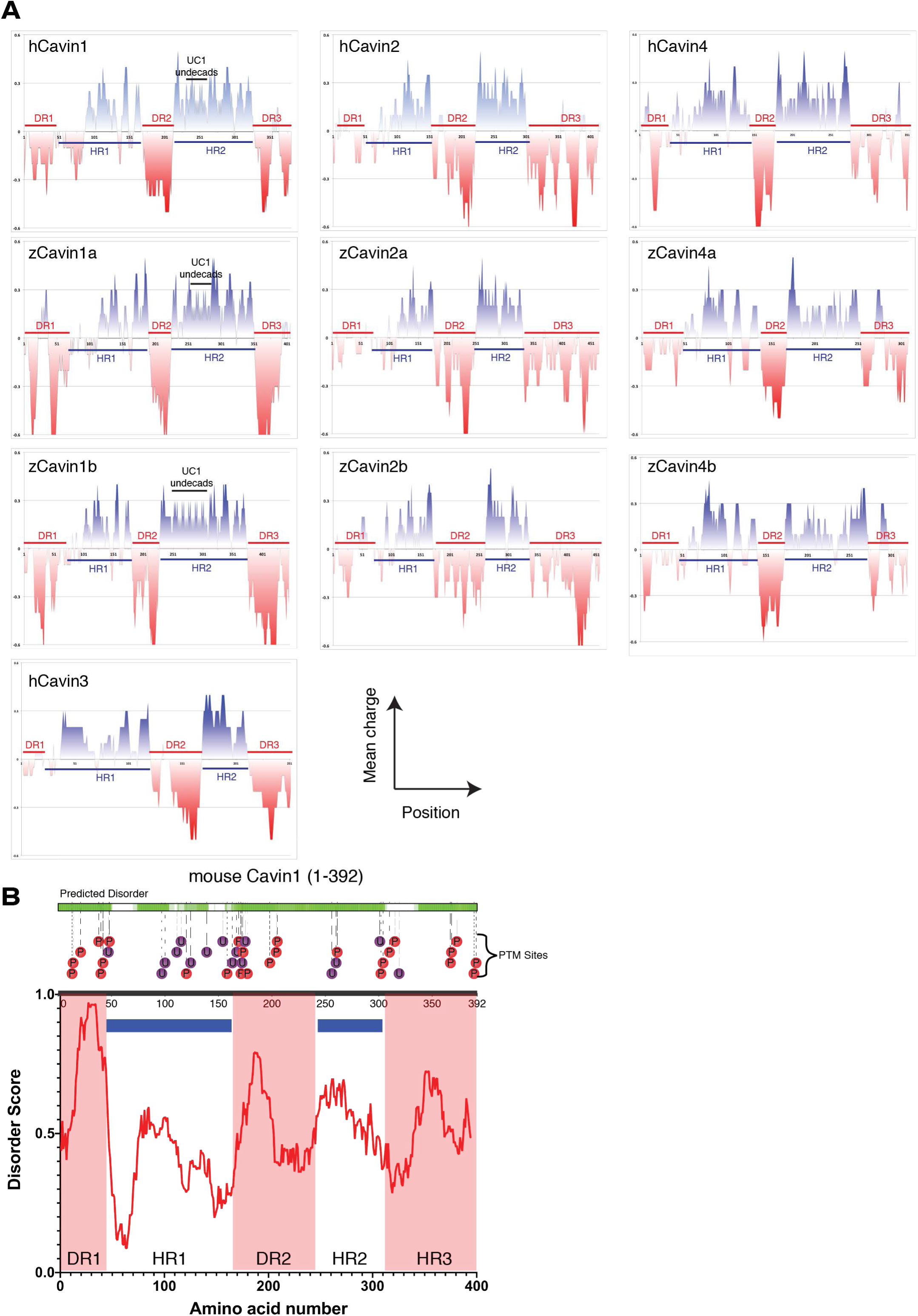
Electrostatic charge distribution and sequence disorder in the Cavin family proteins. **(A)** Protein charge plots of human (h) and zebrafish (z) cavin family proteins performed using the Emboss Server (http://www.bioinformatics.nl/cgi-bin/emboss/charge) (using standard input parameters and a window width of five amino acid residues). **(B)** The Cavin1 sequence was analysed using the D2P2 web server ^15^ for predicted regions of disorder, and also known sites of post-translational modifications.

**Figure S2.**
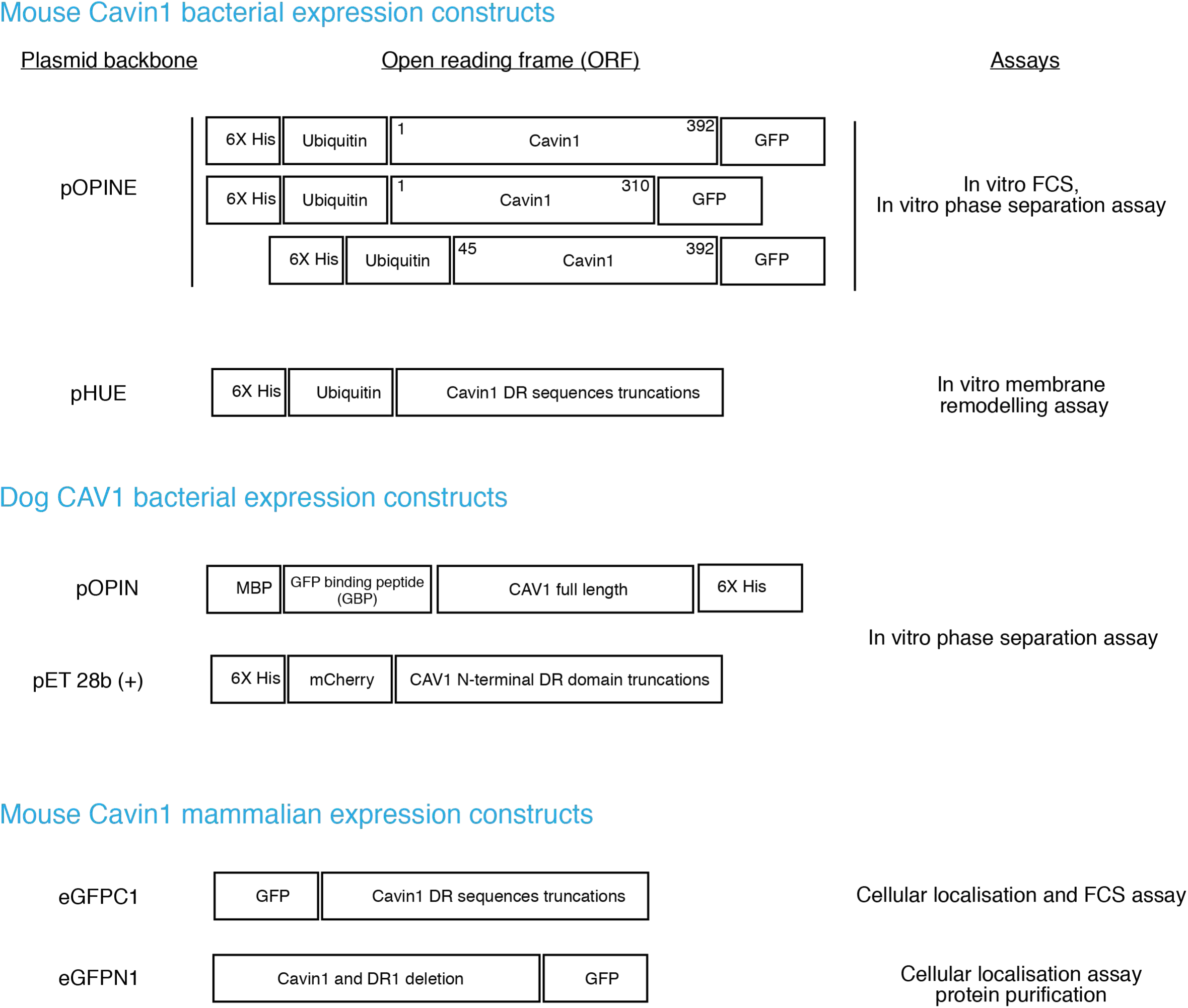
**Schematic representation of protein expression constructs used in this study.**

**Figure S3.**
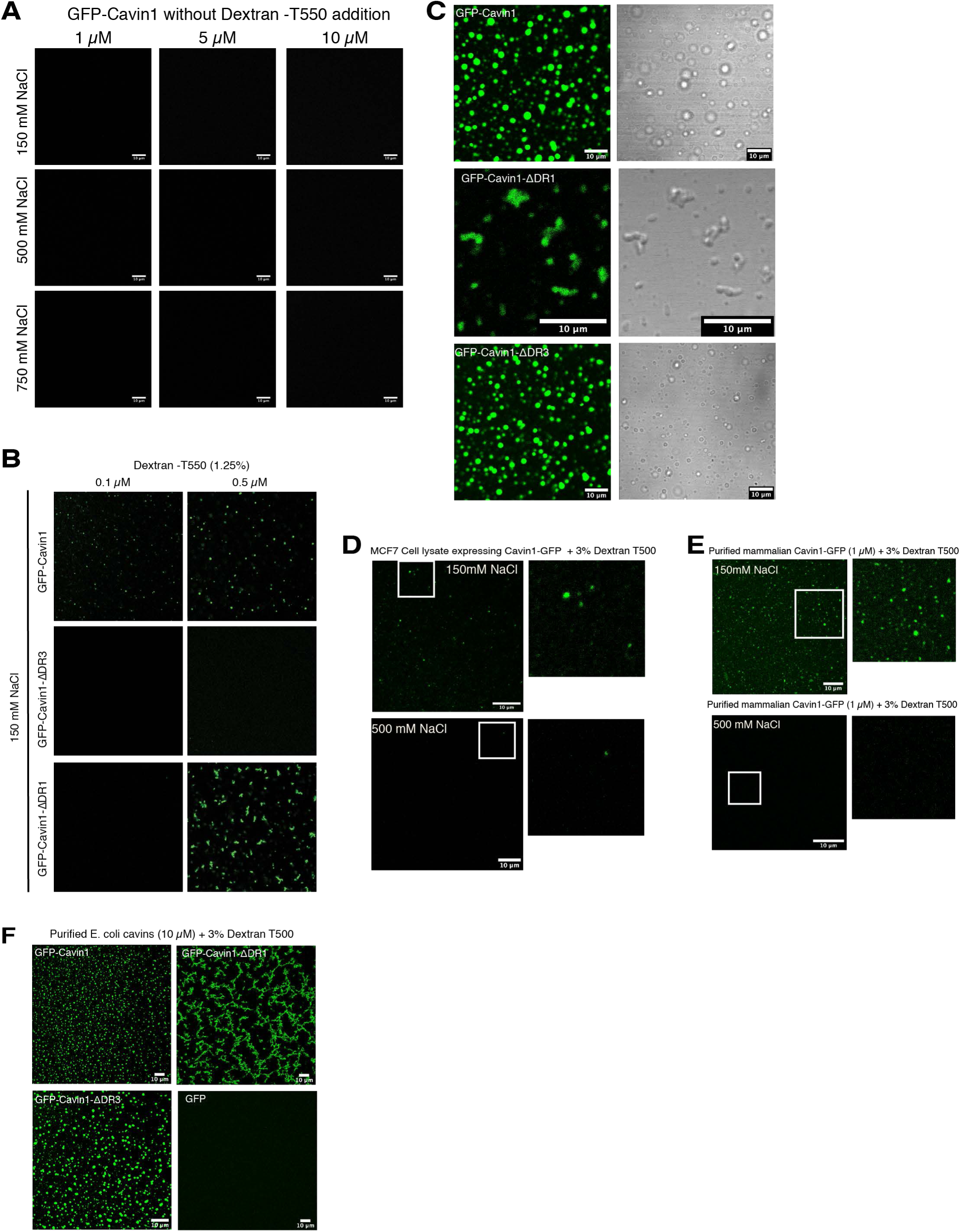
LLPS behaviour of Cavin1 expressed and purified from bacteria and mammalian cells. (**A**) Liquid-liquid phase separation (LLPS) assay with bacterially expressed recombinant Ub- and GFP-tagged Cavin1, at different protein and salt concentrations but in the absence of dextran or other crowding agents. (**B**) At low concentrations, full length Cavin1 still forms liquid droplets, and Cavin1-ΔDR1 still forms coacervates. Cavin1-ΔDR3 is less prone to LLPS at low concentrations compared to the full-length protein. (**C**) LLPS assay performed with GFP tagged Cavin1-ΔDR1 by addition of 1.25% dextran T-500. Fluorescent GFP signal and adjacent bright filed image showing transparent drops unlike non-specific precipitates that are usually non-transparent and milky or brown in appearance. Scale bar – 10 µm. **(D)** LLPS assay performed with Cavin1-GFP expressed and purified from mammalian HEK293 cells. **(E)** MCF7 cell lysates expressing Cavin1-GFP with the addition of 3% dextran T-500 in either 150 mM NaCl or 500 mM NaCl. Scale bar – 10 µm. **(F)** LLPS assay performed with purified E. coli cavins and GFP at higher dextran T-500 concentration (3%).

**Figure S4.**
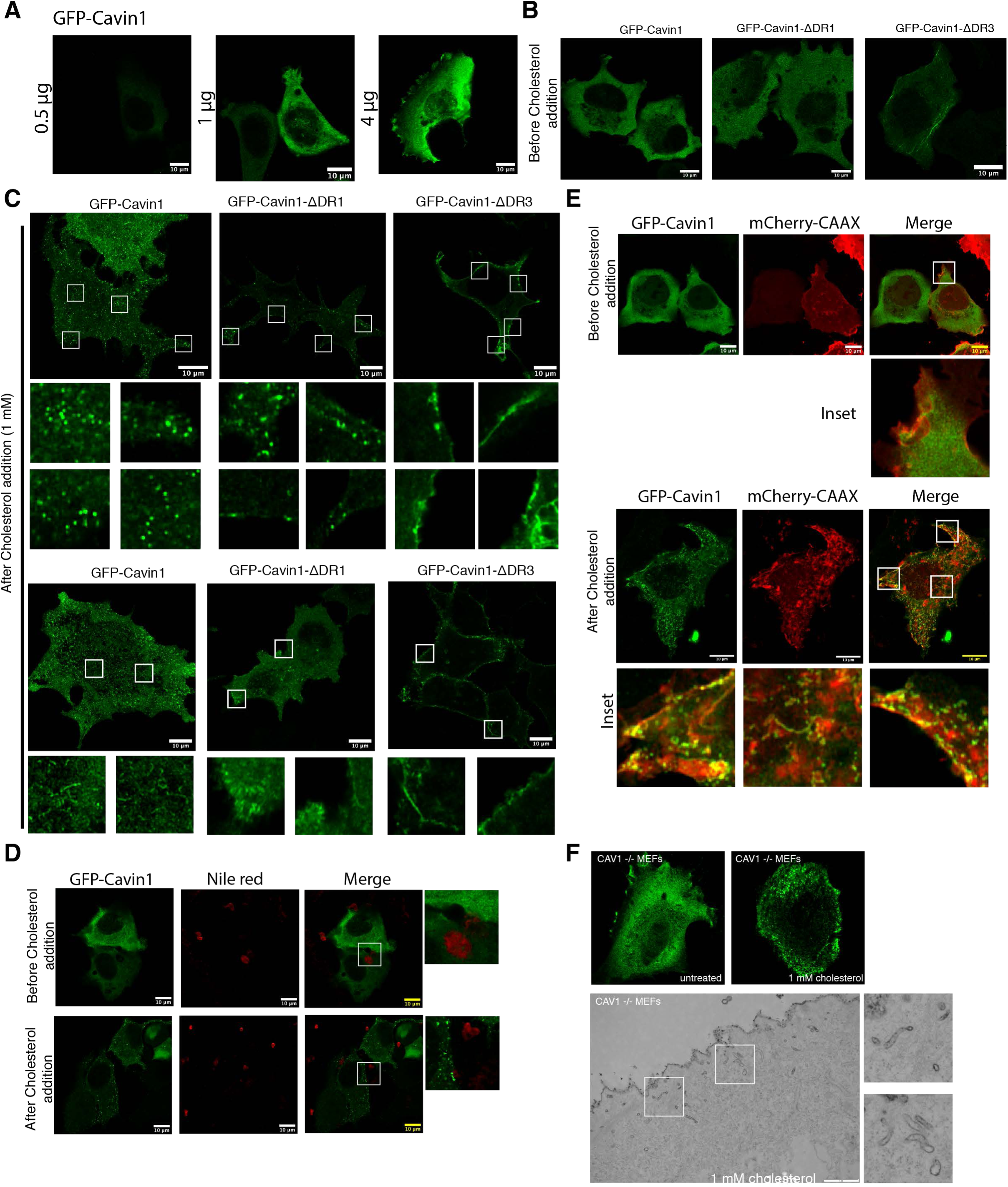
Cavin1 undergoes LLPS and remodels cellular membranes devoid of CAV1. **(A)** MCF7 cells expressing GFP-Cavin1 at varying amount of DNA showing cytosolic distribution. **(B)** MCF7 cells expressing GFP-Cavin1, Cavin1-ΔDR1 showing cytosolic distribution and Cavin1- ΔDR3 showing diffuse localisation (upper panel). **(C)** Addition of a water-soluble form of cholesterol (1 mM added, with effective available cholesterol concentration ∼40 µM) to cells expressing GFP-Cavin1, Cavin1-ΔDR1 and Cavin1-ΔDR3 promotes liquid like droplet formation, membrane recruitment (upper panel) in some cells and tubulation in some cells for GFP-Cavin1 (lower panels). Scale bar – 10 µm. **(D)** MCF7 cells expressing GFP-Cavin1 with cholesterol addition formed GFP-Cavin1 condensates that did not stain with nile red suggesting these structures are not lipid droplets. (**E**) GFP-Cavin1 and mCherry-CAAX co-expression in MCF7 cells before cholesterol addition (upper panel) and after addition of cholesterol (lower panel) showing membrane patches and tubules partially co-localising with mCherry-CAAX. (**F**) CAV1^-/-^ MEF cells expressing GFP-Cavin1 show cytosolic distribution and addition of 1 mM cholesterol causes membrane recruitment of GFP-Cavin1 (left panels) also observed by ruthenium red labelling of membrane surface by EM. Scale bar – 1 µm.

**Figure S5.**
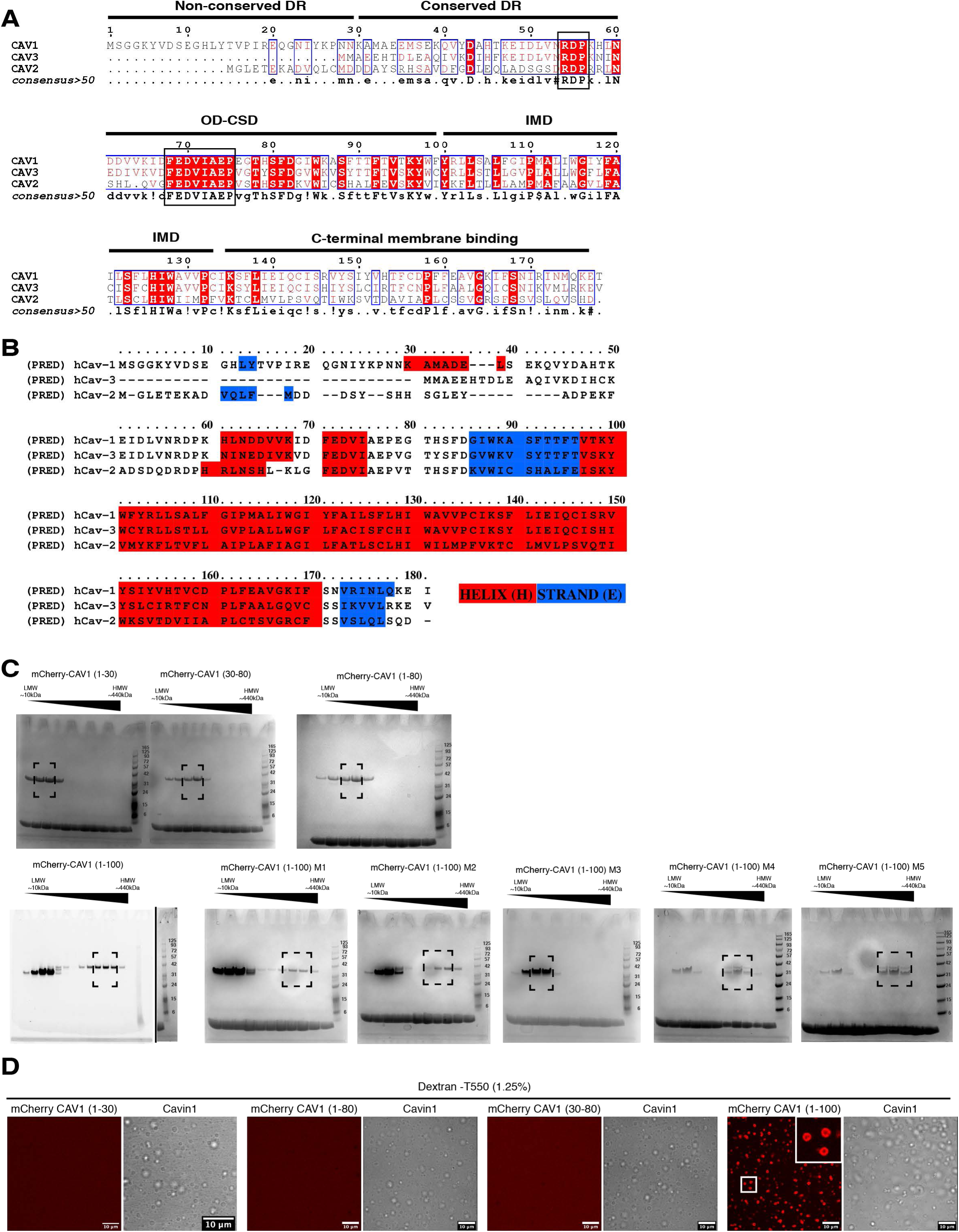
Co-phase separation of CAV1 with Cavin1. **(A)** Amino acid sequence alignment of dog caveolin sequences showing non-conserved and conserved fragments of N-terminal DR region, oligomerization and scaffolding domain (OD-CSD), intramembrane domain (IMD) and C-terminal membrane binding domain. (B) Alignment of human CAV1, CAV2 and CAV3 with secondary structure predictions performed using the Praline webserver (http://www.ibi.vu.nl/programs/pralinewww) ^93^. **(C)** In gel fluorescence images of gel filtration fractions for respective mCherry-tagged CAV1 mutants. **(D)** LLPS assay with mCherry CAV1 (1-30), (30-80), (1-80) and (1-100) and Cavin1. mCherry-CAV1 (1-100) is recruited to cavin1 droplets and undergo LLPS.

**Figure S6.**
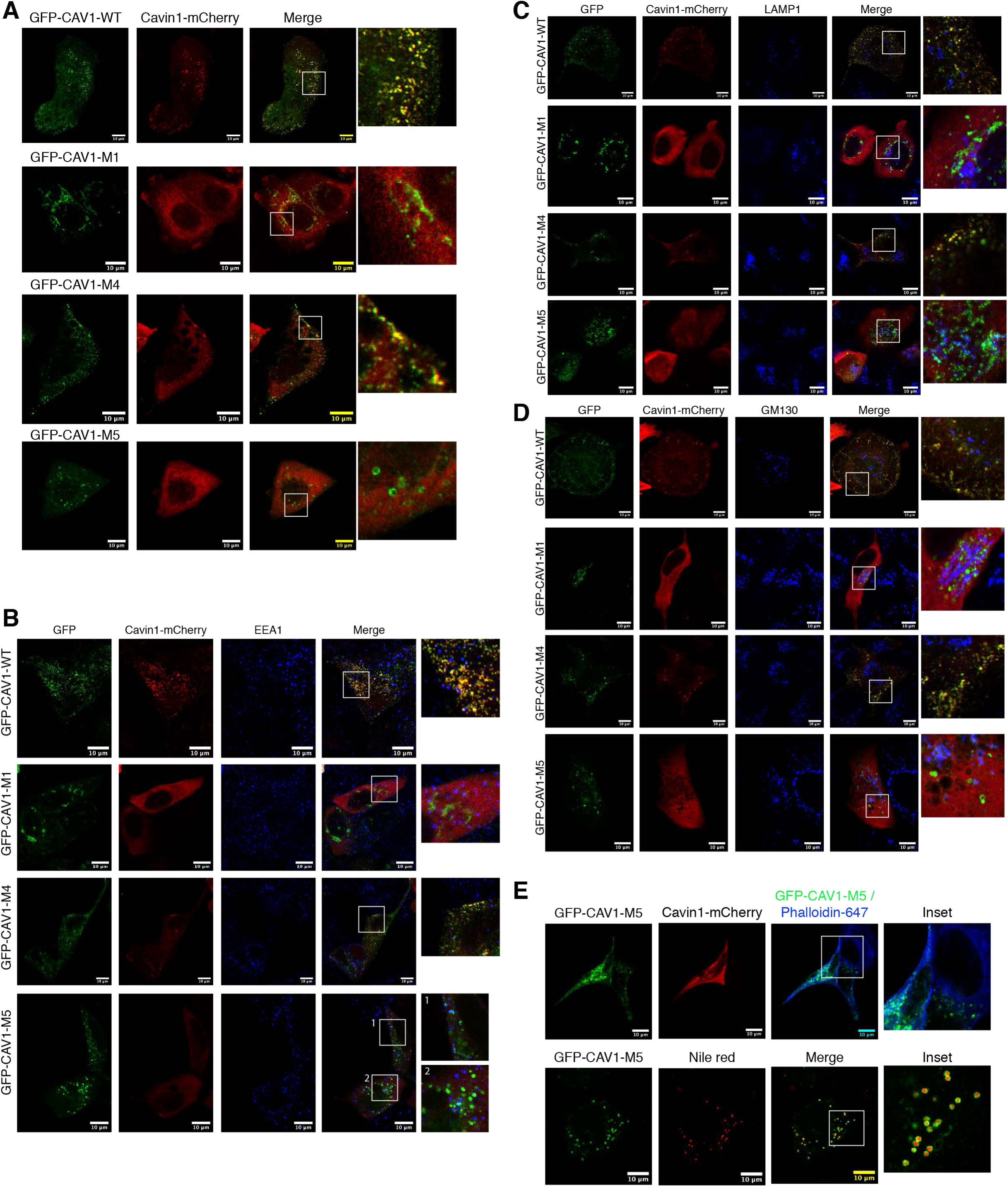
Analysis of GFP-CAV1 mutants co-expressing Cavin1-mCherry in MCF7 cells. GFP tagged CAV1 mutants (green) (Fig. 3) were co-expressed with Cavin1-mCherry in MCF7 cell line **(A)** and fixed cells were immunolabelled for early endosomes (EEA1) **(B)**, lysosomes (LAMP1) **(C)**, golgi membrane (GM130) **(D)**, cellular actin (phalloidin) and nile red (lipid droplets) **(E)** Scale bar – 10 µm

**Figure S7.**
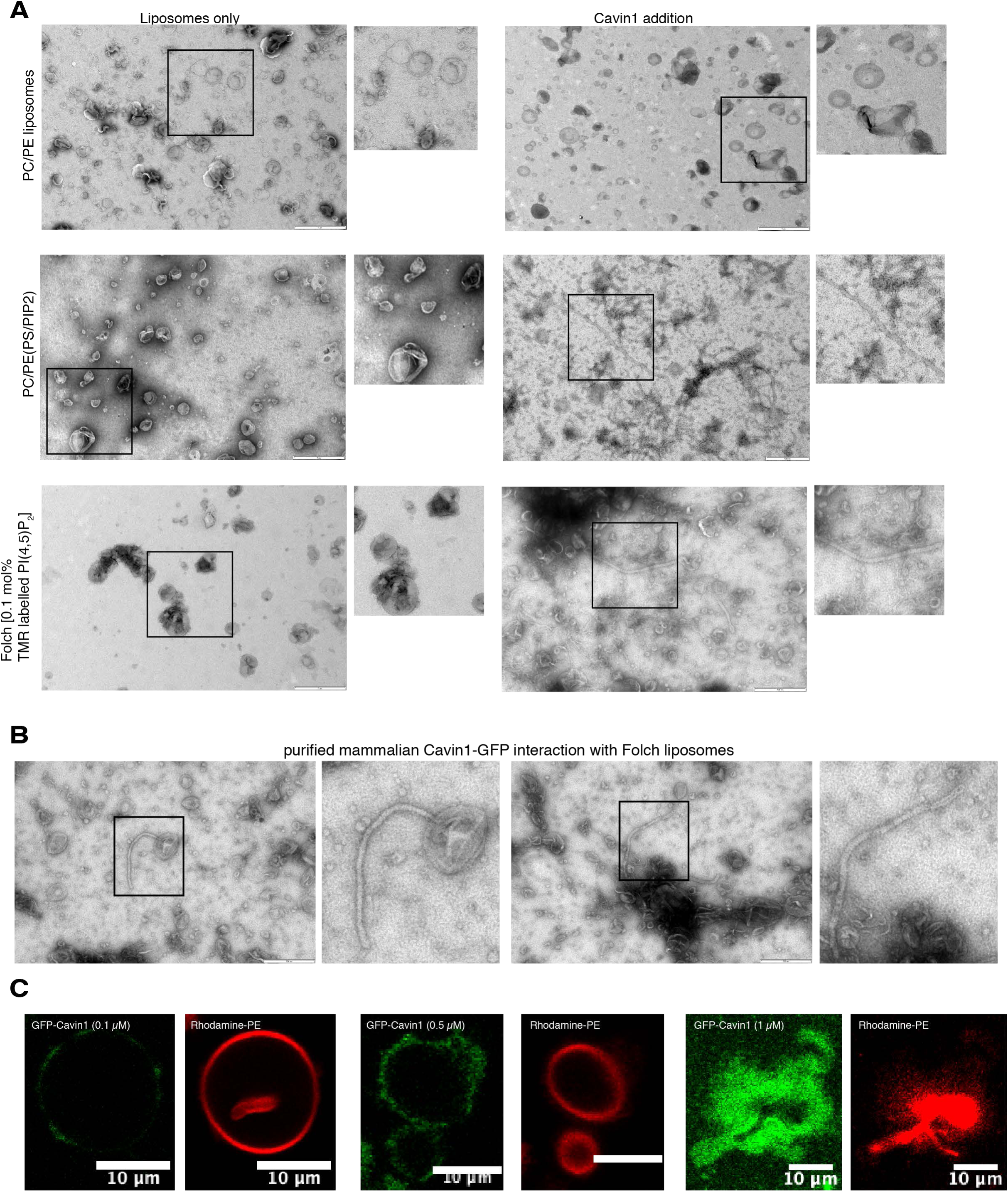
Cavin1 membrane interactions *in vitro*. **(A)** *In vitro* membrane tubulation assay and negative stain electron microscopy was performed after mixing Cavin1 and liposomes consisting of Phosphatidylcholine (PC) and Phosphatidylethanolamine (PE) or PC/PE liposomes containing PI(4,5)P_2_ and Phosphatidylserine (PS) or Folch liposomes containing 0.1 mol% TMR labelled PI(4,5)P_2_ to replicate conditions in Figure 5. (**B**) In vitro membrane tubulation assay performed by mixing mammalian Cavin1-GFP with Folch liposomes, with membrane tubules highlighted in insets. Scale bar – 1 µm. **(C)** Dose dependent GFP-Cavin1 interaction with GMVs containing rhodamine-PE.

**Figure S8.**
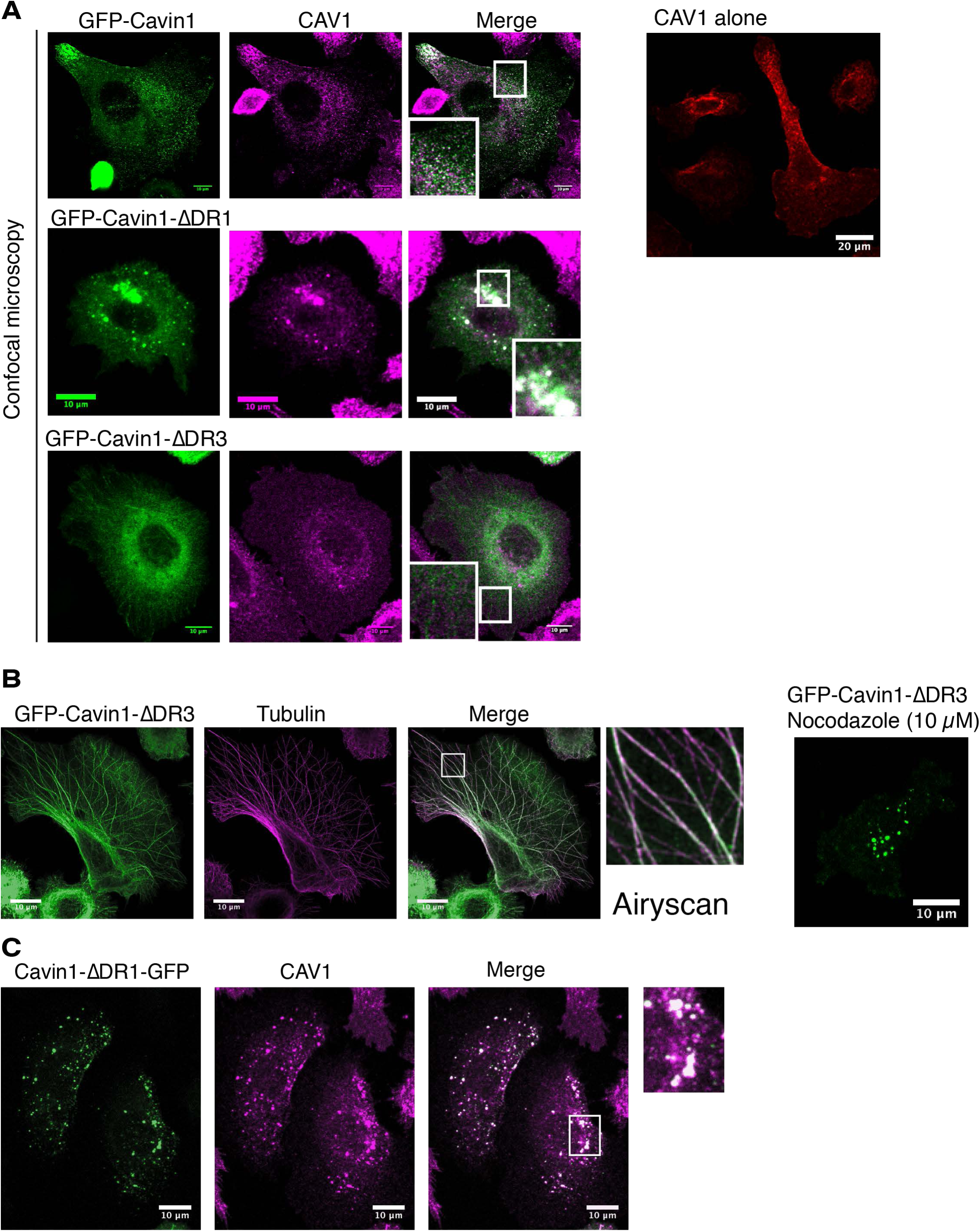
Localisation of Cavin1 with truncated DR1 and DR3 domains. **(A)** Confocal microscopy images of GFP-Cavin1, GFP-Cavin1-ΔDR1 and GFP-Cavin1-ΔDR3 immunolabelled with CAV1 (red) **(B)** GFP-Cavin1-ΔDR3 (green) associates with microtubules (red) in PC3 cells and disperses to the cytosol and forms liquid droplets after nocodazole treatment. Fluorescence images acquired with a Zeiss Airyscan2 microscope. (**C**) Cavin1-ΔDR1-GFP with a C-terminal GFP tag shows a similar intracellular accumulation with CAV1 in PC3 cells as the N-terminal GFP-tagged protein (Fig. 6A), suggesting that the GFP tag does not contribute to this phenotype.

**Figure S9.**
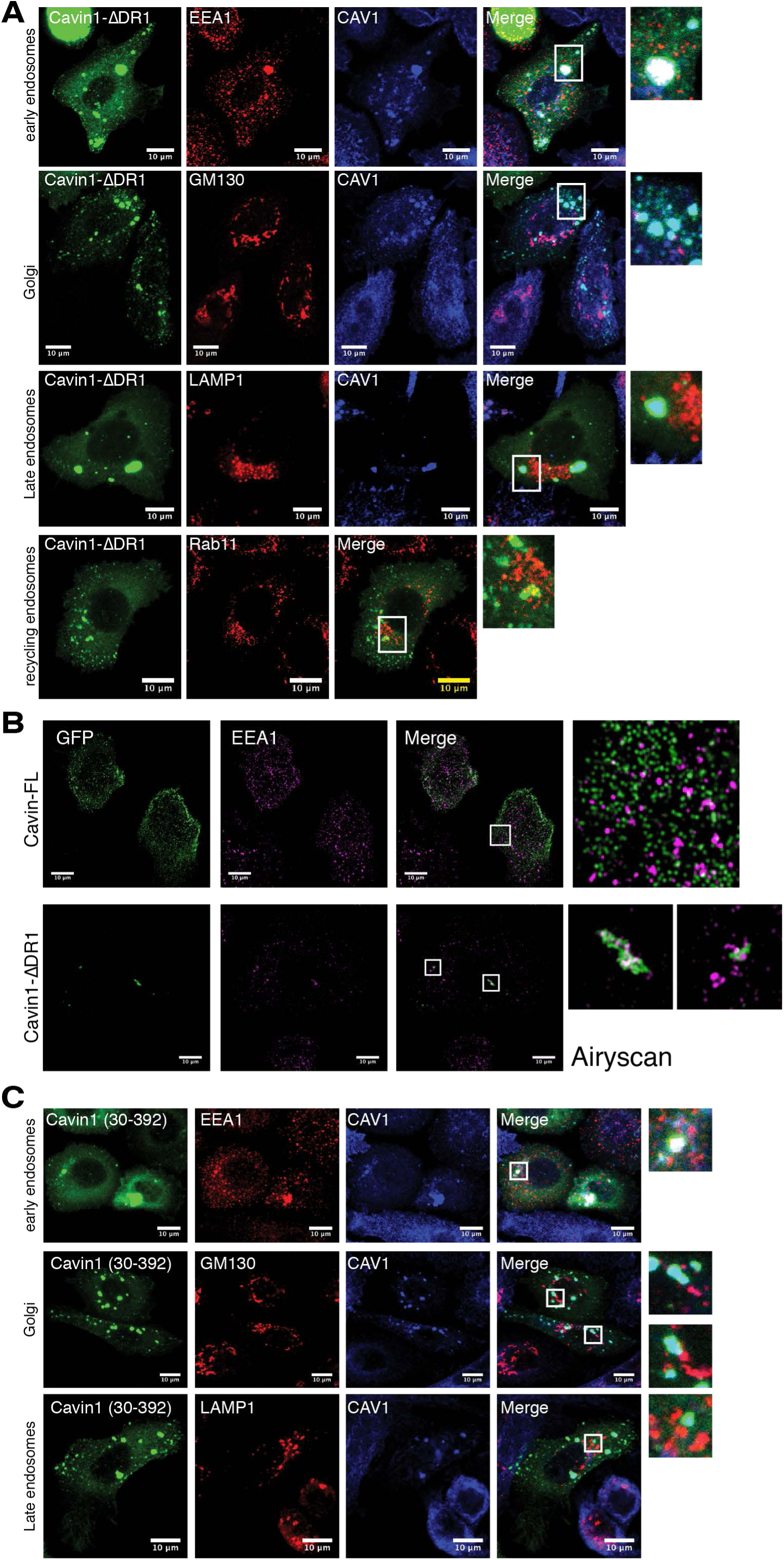
Comparison of Cavin1 truncation mutants with endocytic markers. **(A)** GFP-tagged Cavin1-ΔDR1 (green) was expressed in PC3 cells, and fixed cells were immunolabelled for CAV1 (blue) and different endocytic markers (red) including EEA1, GM130, LAMP1 and Rab11. Only EEA1 showed significant overlap with the internalised Cavin1-ΔDR1 and CAV1 positive structures. **(B)** High-resolution images of GFP-tagged Cavin1 and Cavin1-ΔDR1 (green) in PC3 cells compared with EEA1 (magenta) acquired with a Zeiss Airyscan2 microscope. (**C**) As for (**A**) but cells expressing GFP-tagged Cavin1(30-392). Cavin1(30-392) accumulates at intracellular structures with CAV1 and positive for EEA1 labelling similarly to Cavin1-ΔDR1 with the full deletion of the DR1 domain.

**Figure S10.**
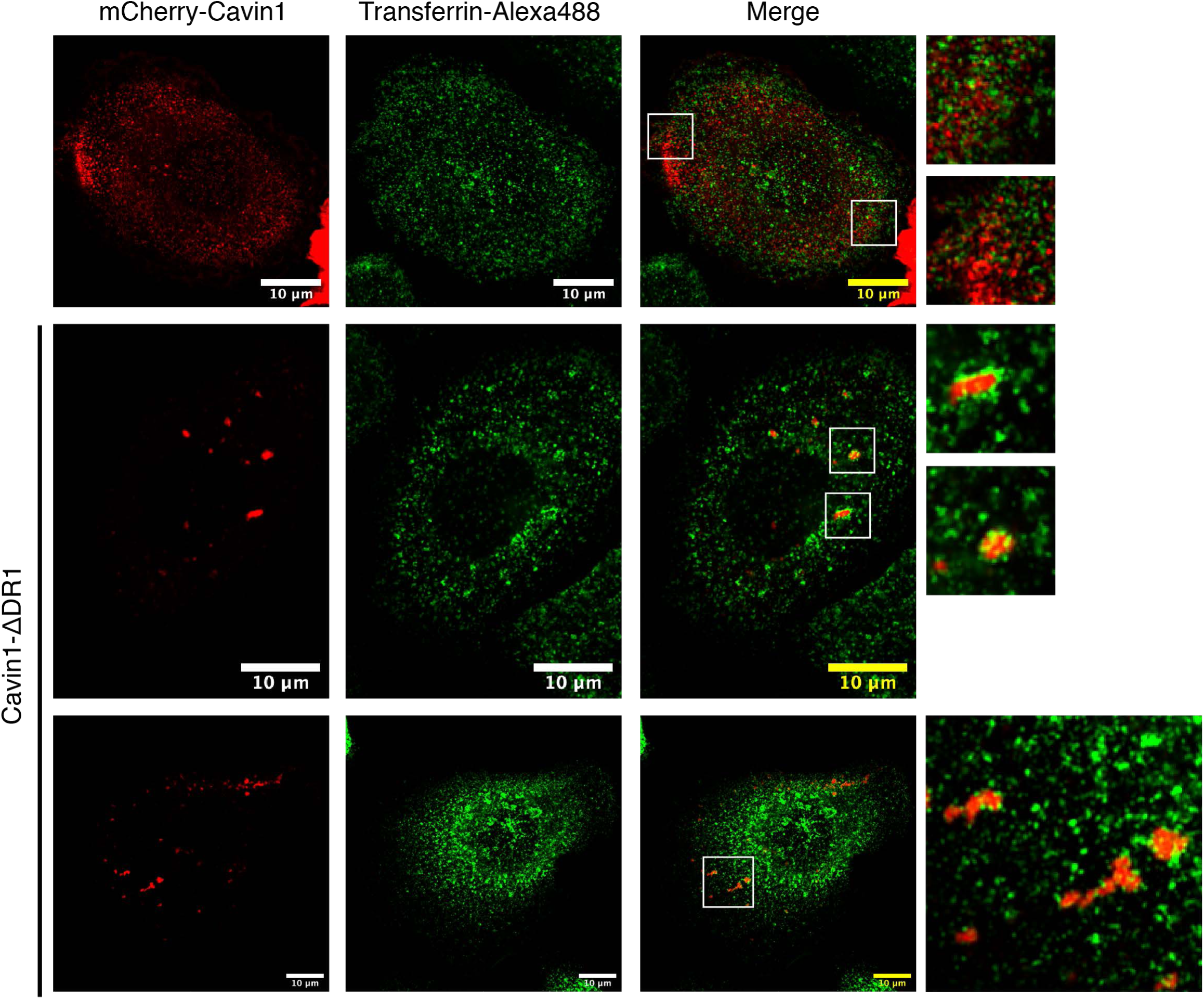
Cavin1-ΔDR1 shows colocalisation with internalised transferrin. Transferrin uptake assay was performed in PC3 cells expressing either mCherry-tagged Cavin1 or Cavin1-ΔDR1 (red) with transferrin Alexa-488 (green). Wild-type mCherry-Cavin1 showed no colocalization with endocytosed transferrin whereas mCherry-Cavin1-ΔDR1 formed large structures (red) with transferrin positive endosomes surrounding them.

**Figure S11.**
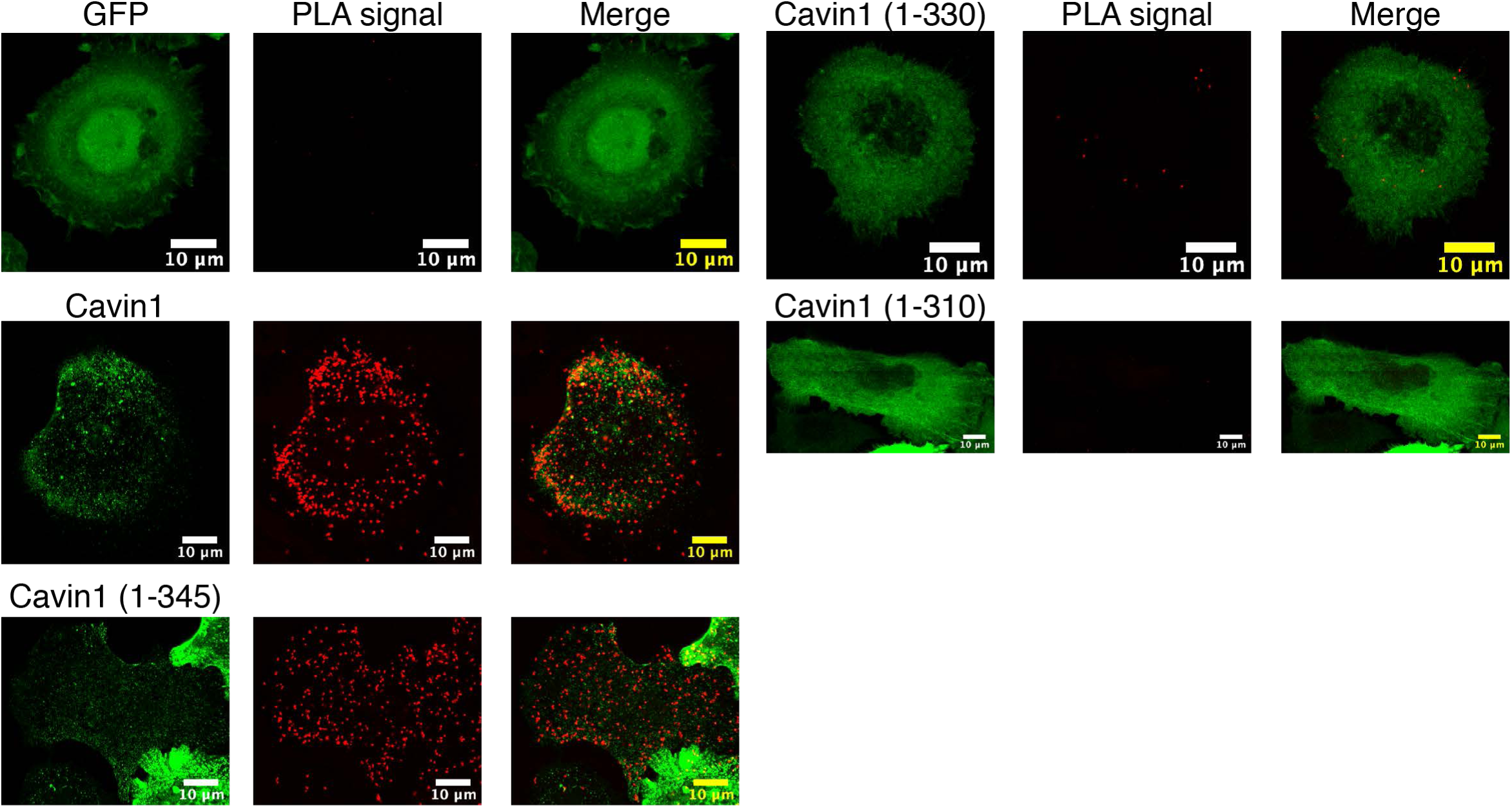
PLA assay of Cavin1 interactions with CAV1. Representative images of proximity ligation assays of Cavin1 and CAV1 interactions, with GFP-tagged Cavin1 mutants in green and PLA signal in red. Scale bar – 10 μm. Related to Fig. 7C.

**Figure S12.**
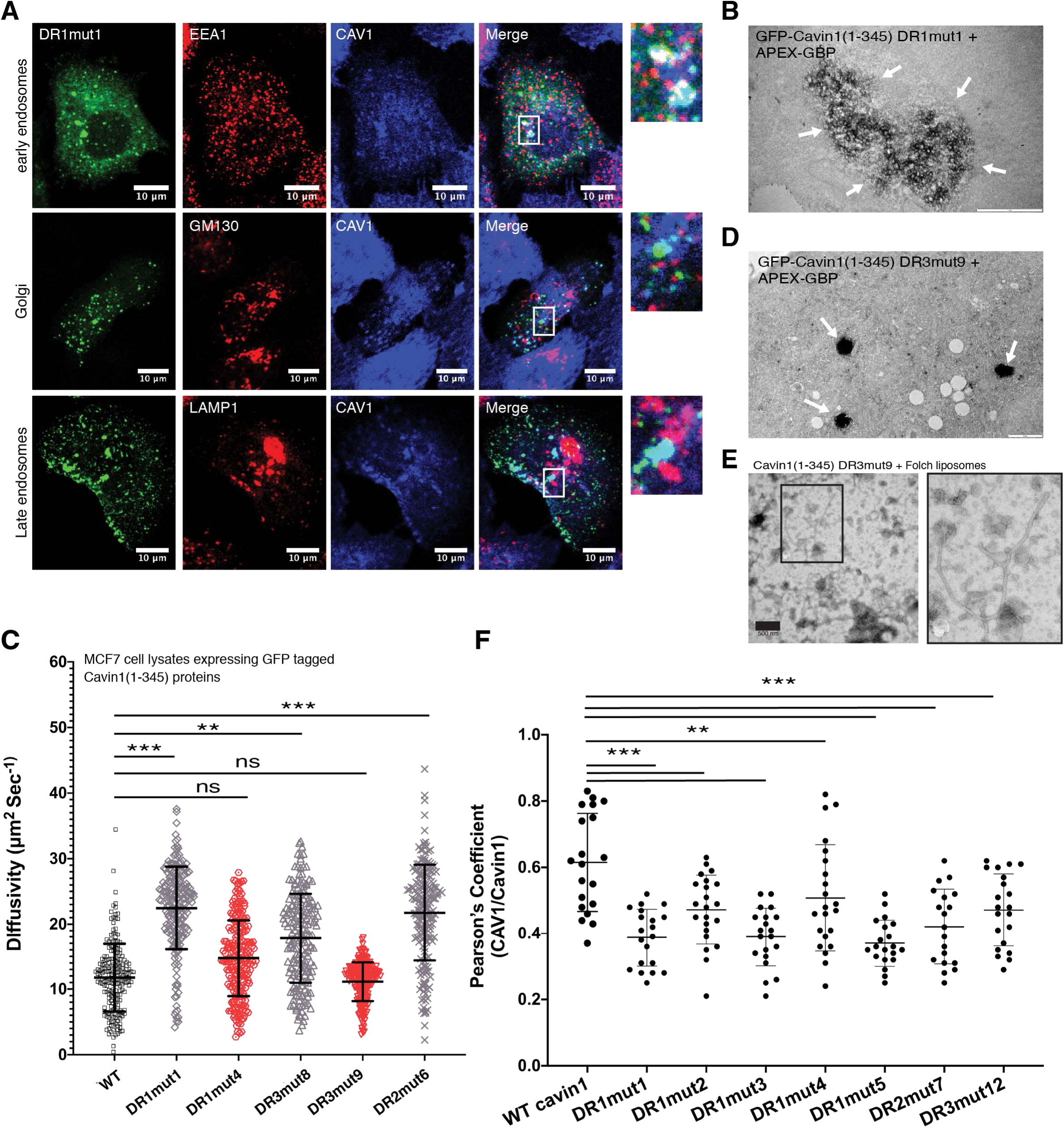
Localisation and membrane remodelling by Cavin1(1-345) mutant proteins. **(A)** GFP-tagged Cavin1(1-345) mutant DR1mut1 was expressed in PC3 cells, and fixed cells were immunolabelled for CAV1 (blue) and different endocytic markers (red) including EEA1, GM130, and LAMP1. Like the complete deletion of the residues 1-30 in the Cavin1 DR1 region (**Fig. S9C**) Cavin1(1-345) mutant DR1mut1 shows significant overlap with CAV1 and EEA1 positive internal structures. **(B)** APEX-GBP labelling of GFP-tagged Cavin1(1-345) mutant DR1mut1 shows accumulation and clustering with internal membrane vesicles (arrows). (**C**) The diffusion rate measured by FCS of selected GFP-tagged Cavin1(1-345) DR mutants in lysates after expression in MCF7 cells (lacking endogenous Cavins and Caveolins). N = 3, n = 15-25. Error bars indicate mean ± SD, **P<0.05, *** P<0.001, ns – not significant. (**D**) APEX-GBP labelling of GFP-tagged Cavin1(1-345) mutant DR3mut9 shows droplet localisation (arrows). (**E**) Purified Ub-tagged Cavin1(1-345) mutant DR3mut9 was mixed with unilamellar Folch liposomes (extruded to 400 nm diameter) and analysed by negative stain EM (1% uranyl acetate). This mutant is able to remodel and tubulate these synthetic membranes, although with a slightly larger diameter than wild-type Cavin or Cavin1(1-345) (Fig. 4D). **(F)** GFP-Cavin1 and various DR mutants of Cavin1 (1-345) were expressed in PC3 cell line and immunolabelled for CAV1 after fixation. The co-colocalization of GFP tagged cavin variants and CAV1 was quantified by Pearson’s correlation coefficient. N = 2, n = 8-12. Error bars indicate mean ± SD. **P<0.05, *** P<0.001.

**Figure S13.**
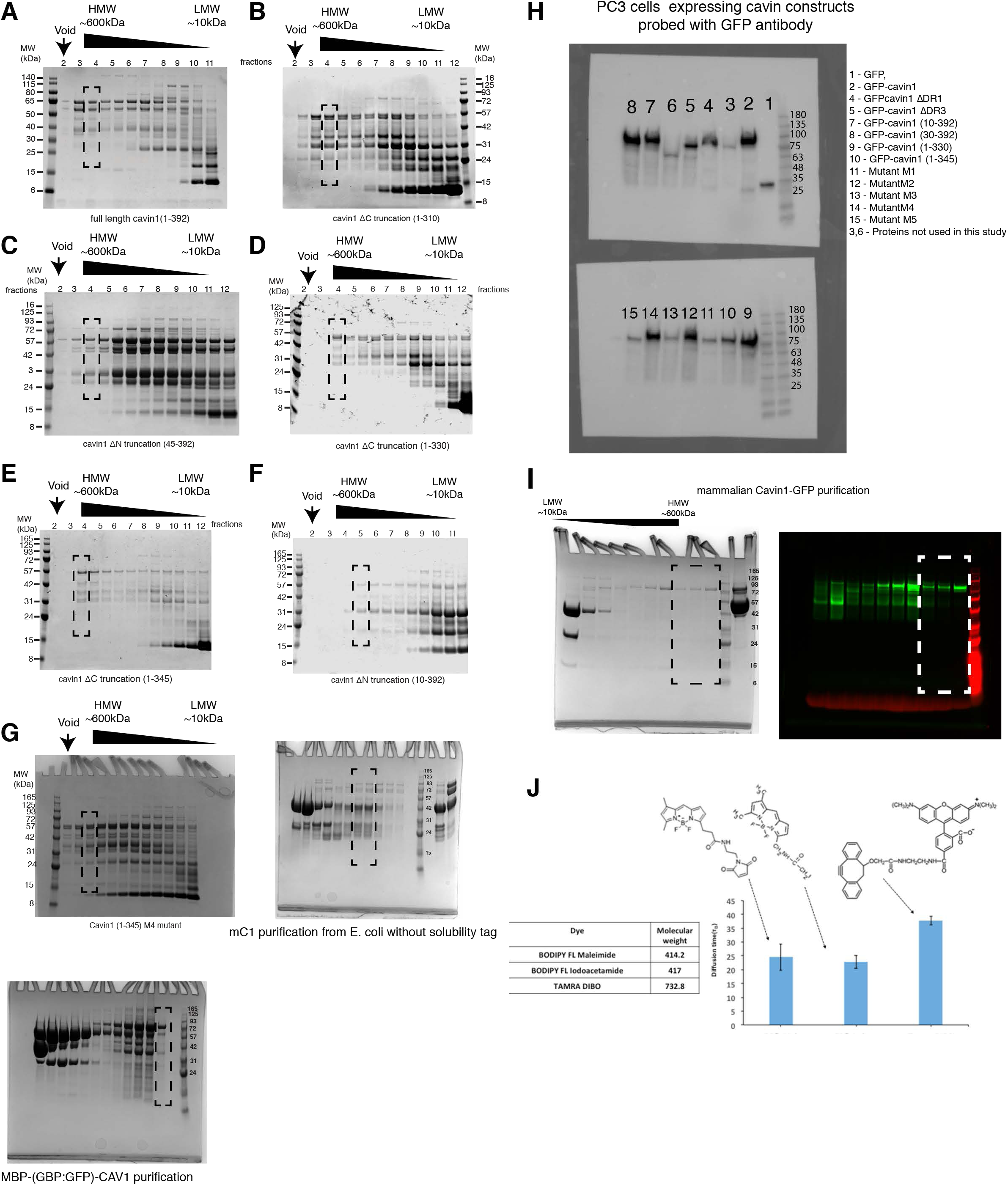
(**A to G**) Gels showing purified recombinant Cavin1 proteins used in this study. (**H**) Western blot showing expression of GFP tagged mutants expressed in PC3 cell line probed with anti-GFP antibody. (**I**) SDS-PAGE and in gel fluorescence profile of Cavin1-GFP purified from HEK cells using GFP nanobody and subjected size exclusion chromatography on superose 6 (10/300) column. (**J**) The diffusion time measurements for three dyes performed before each FCS session.

**Movie S1. (related to Fig. 4B).**

Cryoelectron tomography (CryoET) of Cavin1-coated membrane tubules. The movies shows a series of images panning through the three-dimensional tomographic volume. Striated protein densities are observed coating the relatively heterogeneous membrane tubules.

**Movie S2. (related to Fig. 6B).**

GFP-tagged Cavin1-ΔDR3 was expressed in PC3 cells and photobleaching was performed on a small region along microtubules coated with GFP tagged mutant protein. Images were acquired one frame per second.

**Movie S3. (related to Fig. 6C).**

GFP-tagged Cavin1-ΔDR3 was expressed in PC3 cells and treated with nocodazole (10 µM). Photobleaching was performed on a small region containing liquid droplets of mutant protein and images were acquired one frame per second.

**Movie S4. (related to Fig. 6F).**

GFP-tagged Cavin1 and Rab5a-mCherry were co-expressed in PC3 cells and images were acquired one frame per four seconds.

**Movie S5. (related to Fig. 6F).**

GFP-tagged Cavin1-ΔDR1 and Rab5a-mCherry were co-expressed in PC3 cells and images were acquired one frame per four seconds.

**Movie S6. (Related to Fig. 8E).**

GFP-Cavin1 DR3mut9 mutant expressed in PC3 cell line and photobleaching was performed on protein droplets dispersed in cytosol. Images were acquired one frame per two seconds.

